# SAMHD1 controls innate immunity by regulating condensation of immunogenic self RNA

**DOI:** 10.1101/2022.07.12.499661

**Authors:** Shovamayee Maharana, Stefanie Kretschmer, Susan Hunger, Xiao Yan, David Kuster, Sofia Traikov, Thomas Zillinger, Marc Gentzel, Nagaraja Chappidi, Nadja Lucas, Katharina Isabell Maser, Henrike Maatz, Alexander Rapp, Virginie Marchand, K. Venkatesan Iyer, Akshita Chhabra, Young-Tae Chang, Yuri Motorin, Norbert Hubner, Gunther Hartmann, Anthony Hyman, Simon Alberti, Min Ae Lee-Kirsch

## Abstract

Recognition of pathogen-derived foreign nucleic acids is central to innate immune defense. This requires discrimination between structurally highly similar self and nonself nucleic acids to avoid aberrant inflammatory responses as in the autoinflammatory disorder Aicardi-Goutières syndrome (AGS). How vast amounts of self RNA are shielded from immune recognition to prevent autoinflammation is not fully understood. Here we show that SAM domain and HD domain-containing protein 1 (*SAMHD1)*, one of the AGS-causing genes, functions as a single-stranded RNA (ssRNA) 3’exonuclease, the lack of which causes cellular RNA accumulation. Increased ssRNA in cells leads to dissolution of RNA-protein condensates, which sequester immunogenic double-stranded RNA (dsRNA). Release of sequestered dsRNA from condensates triggers activation of antiviral type I interferon via retinoic acid-inducible gene I-like receptors. Our results establish SAMHD1 as a key regulator of cellular RNA homeostasis and demonstrate that buffering of immunogenic self RNA by condensates regulates innate immune responses.

## Introduction

In view of the coronavirus pandemic, understanding innate immune responses as the first line of defense against viral infections has gained importance in an unprecedented way. Detection of viruses occurs primarily through recognition of viral nucleic acids by a limited number of pattern recognition receptors. Cytosolic foreign RNA is detected by retinoic acid-inducible gene I (RIG-I) and melanoma differentiation-associated gene 5 (MDA5) (Kato et al., 2006; Kawai et al., 2005), which upon engagement activate antiviral type I interferon (IFN) (Schlee and Hartmann, 2016). In addition, there are several RNA receptors that require additional licensing signals such as type I IFN to initiate antiviral signalling, including dsRNA-activated protein kinase R (PKR), IFN-induced protein with tetratricopeptide repeats 1 (IFIT1), and 2′-5′-oligoadenylate synthetase 1 (OAS1) (Schlee and Hartmann, 2016). Targeted detection of foreign RNA within the cytosol, an environment containing vast amounts of highly diverse endogenous RNA species that share similar properties with viral RNA, remains imperfect, posing a considerable challenge to the innate immune system. While our understanding of cytoplasmic RNA receptors has increased substantially in recent years, how these receptors faithfully spot a few copies of viral RNA amidst a large number of endogenous RNA molecules is still imperfectly understood.

As erroneous recognition of self nucleic acids can cause autoinflammation, cells must be able to prevent inappropriate innate immune activation. One way to distinguish self from nonself RNA is by detecting structural properties of foreign RNA, such as extended double-stranded regions (Schlee and Hartmann, 2016). Such regions are also present in endogenous RNAs derived from inverted Alu repeats, but these are modified by adenosine deaminase (ADAR), thereby preventing recognition by the dsRNA sensor MDA5 (Liddicoat et al., 2015). Self RNA is also rendered invisible to the immune system by adding a cap structure to the 5’ end. Foreign RNAs lack a cap but often contain triphosphate or diphosphate at the 5′ end, which are detected by RIG-I (Goubau et al., 2014; Hornung et al., 2006; Pichlmair et al., 2006). Discrimination of self from nonself RNA is further achieved by confinement of RNA sensors such as endosomal Toll-like receptor (TLR) 7 or TLR8 to membrane-enclosed organelles (Pelka et al., 2016). However, cells also harbour many membraneless compartments that contain large amounts of RNA (Banani et al., 2017). Membraneless compartments such as nucleoli or stress granules are also referred to as condensates because they assemble from proteins and RNA by liquid-liquid phase separation (Banani et al., 2017). But whether the ability of condensates to sequester RNA is involved in immune recognition has not been investigated so far.

Aberrant sensing of self nucleic acids due to mutations in enzymes involved in nucleic acid metabolism underlies Aicardi-Goutières syndrome (AGS), an autoinflammatory disorder characterized by constitutive type I IFN activation (Lee-Kirsch, 2017; Uggenti et al., 2019). One of the AGS-causing genes, *SAMHD1*, functions as a deoxynucleotide (dNTP)-degrading triphosphohydrolase, which regulates cellular dNTP pools (Goldstone et al., 2011; Powell et al., 2011; Rice et al., 2009) and is implicated in HIV-1 restriction and in genome stability (Hrecka et al., 2011; Kretschmer et al., 2015; Laguette et al., 2011; Lahouassa et al., 2012). SAMHD1 was also shown to promote DNA end resection at stalled replication forks and DNA double-strand breaks independent of its dNTPase activity (Coquel et al., 2018; Daddacha et al., 2017). In addition, SAMHD1 is suggested to possess ribonuclease (RNase) activity *in vitro* (Beloglazova et al., 2013). However, the physiological role of its RNase function and its possible role in aberrant innate immune activation has remained unclear.

In our study, we investigated the RNA-associated functions of SAMHD1 and establish that SAMHD1 acts as an ssRNA-degrading RNase in cells. Loss of SAMHD1 causes cellular accumulation of RNA leading to impaired formation of RNA-containing condensates by liquid-liquid phase separation. We demonstrate that condensates normally sequester immunostimulatory dsRNA, thereby preventing innate immune recognition of self RNA, but that dsRNA is released from condensates in AGS patient cells, causing aberrant activation of the innate immune system. Thus, maintaining self RNA tolerance by regulating the spatial organisation of RNA through phase separation is an essential cellular function of SAMHD1.

## Results

### SAMHD1 functions as RNase *in vitro* and *in cellulo*

To investigate the RNase activity of SAMHD1 (**Figure S1A**), we purified recombinant human GST-SAMHD1 (**Figure S1B**). Previous studies have shown that SAMHD1 is functional in a tetrameric form (Yan et al., 2013). To characterise the functionality of purified SAMHD1, we performed size exclusion chromatography, and found that GST-SAMHD1 was monomeric, but oligomerized in the presence of 200 µM dGTP (**Figure S1C**), consistent with a dGTP-dependent mechanism of oligomerisation (Ji et al., 2013). We next assessed the purity of recombinant SAMHD1 by mass spectrometric analysis, which revealed absence of contaminant nucleases in the preparation (**Table S1**). We then performed *in vitro* nuclease assays using purified recombinant human GST-SAMHD1 and various RNA substrates. In the presence of Ca^2+^, wild type SAMHD1 efficiently degraded total RNA, poly(A)-mRNA as well as single-stranded RNA (ssRNA) oligonucleotides with a 5’-cap or 5’-triphosphate, respectively (**Figure 1A**). In contrast, two AGS-associated SAMHD1 mutants (Q548X and H167Y) did not show any RNase activity (**Figure 1A**). The RNase activity of the phosphorylation mutant T592A (**Figure S1D**) did not differ from wild type SAMHD1 (**Figure 1A**), suggesting that phosphorylation at T592 is not required for RNase activity and only relevant for regulating the dNTPase activity of SAMHD1 (White et al., 2013). SAMHD1 did neither degrade ssRNA with a 3’-monophosphate nor double-stranded RNA (dsRNA), small interfering RNA (siRNA), RNA:DNA hybrids, or RNA:protein complex 70S ribosome, while it only partially degraded primary micro-RNA consisting of ssRNA with double-stranded stem loops (**Figure 1A; Figure S1H**), indicating that SAMHD1 acts as a 3’-ssRNA exonuclease. Neither oligomerisation nor phosphorylation of SAMHD1 interfered with RNase activity, which was dependent on divalent cations (**Figures S1C-S1G**).

**Figure 1.**
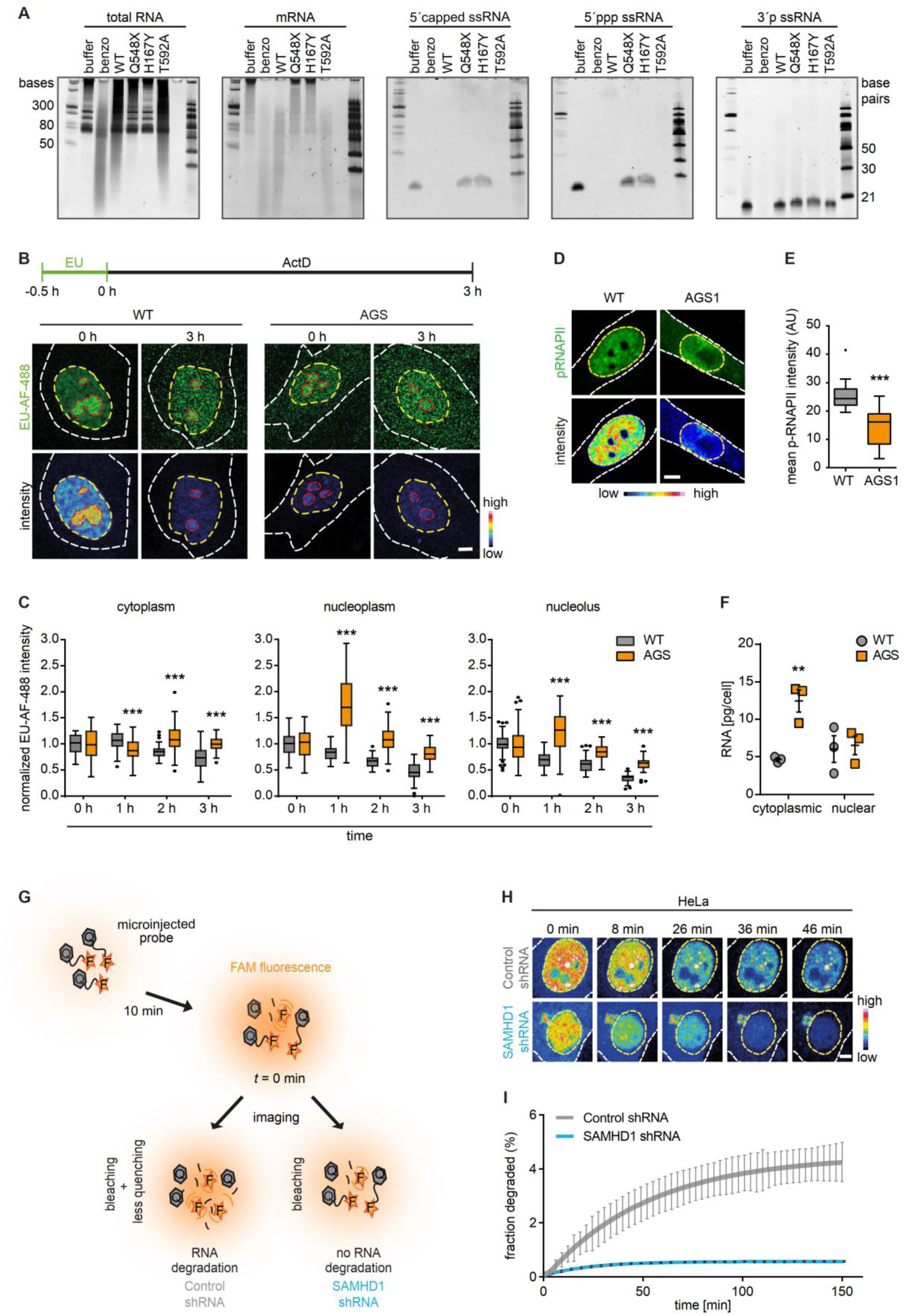
SAMHD1 functions as RNase *in vitro* and *in cellulo*. (**A**) RNase activity of wild type (WT) and mutant (Q548X, H167Y, T592A) SAMHD1 towards indicated substrates. Buffer, negative control. Benzo, benzonase, positive control. First lane, ssRNA ladder; last lane, dsRNA ladder. (**B**) EU-Alexa Fluor 488 (green) and color-coded intensity of metabolically labeled nascent RNA in wild type (WT) and AGS fibroblasts. Scale bar, 5 μm. (**C**) Quantification of normalized RNA intensity. n = 51-82 cells. Box plots: center line, median; box, interquartile range; whiskers, 1.5x interquartile range; dots, outliers. Two-way ANOVA, ***, p<0.001. (**D**) Representative images of immune-stained phosphorylated RNA pol II, (pRNAPII; green) along with color-coded intensity in wild-type (WT) and AGS1 fibroblasts. Outlines of cell and nucleus are marked with white and yellow dashed lines, respectively. Scale bar, 5 μm. (**E**) Quantification of transcriptionally active RNA pol II. n = 11-19 cells. Box plots: center line, median; box, interquartile range; whiskers, 1.5x interquartile range; dots, outliers. Mann-Whitney U test. ***, p<0.001. (**F**) Total RNA levels in cytosol and nuclei from three different passages of AGS1 patient cells and three different wild type controls. Two-way ANOVA with Sidak’s multiple comparisons test. **, p<0.01. (**G**) Schematic of *in cellulo* RNA degradation assay. A 22 nt ssRNA probe labeled with 5’ FAM (fluorescence reporter, F) and 3’ TAMRA (quencher, Q) was microinjected into nuclei of HeLa cells expressing either control-shRNA or SAMHD1-shRNA and imaged after 10 min. (**H**) Representative images of intensity-coded FAM fluorescence at indicated time points of time series starting 10 min (*t* = 0) after microinjection of FAM-TAMRA-labeled ssRNA oligonucleotide. Scale bar, 5 μm. (**I**) Quantification of FAM fluorescence in nuclei over time, where *t* = 0 is 10 min after microinjection of ssRNA probe. (**A-I**) Data are representative of at least three independent experiments.

Given the lack of RNase activity of AGS-associated SAMHD1 mutants *in vitro*, we hypothesized that cells carrying these mutations should show defects in RNA degradation. We examined global RNA degradation in fibroblasts from an AGS patient (AGS1; compound heterozygous for R290H and Q548X) by metabolic labelling of RNA with 5-ethynyl uridine (EU) for 30 min followed by inhibition of new transcription by actinomycin D and ligation of Alexa Fluor (AF)-488 to incorporated EU nucleosides. Global transcription in SAMHD1-deficient fibroblasts was lower compared to control cells as indicated by the low EU-Cy5 intensity at *t* = 0 and the low level of transcriptionally active RNAPII in AGS cells (**Figure 1B; Figures 1D and 1E**). In agreement with the nuclear localization of SAMHD1 (Kretschmer et al., 2015), we observed a higher fraction of EU-labelled RNA in both the nucleolus and nucleoplasm of AGS patient cells after 3 hours as compared to wild type fibroblasts (**Figures 1B and 1C**). Remarkably, we also noted a higher fraction of EU-labeled RNA in the cytoplasm of SAMHD1-deficient fibroblasts that was not seen in wild type cells (**Figure 1B and 1C**), suggesting that higher nuclear RNA levels lead to a concomitant increase in RNA concentration in the cytosol presumably by RNA transport or leakage out of the nucleus. In line with the findings observed by pulse metabolic labelling of RNA, total RNA levels measured by spectrophotometry, which reflect the equilibrium state, were higher in the cytosol of AGS patient cells compared to controls (**Figure 1F**).

To test if the higher RNA levels in AGS patient cells were indeed a direct effect of a lack of SAMHD1 RNase activity, we followed degradation of a 22 nt ssRNA oligonucleotide reporter RNA microinjected into cells. This RNA was dual-labelled with a 5’ fluorophore (FAM) and a 3’ quencher (TAMRA) and fluoresces upon nucleolytic degradation of the 3’ end. HeLa cells were microinjected with the RNA probe and the degradation rate of the RNA probe was computed from the FAM fluorescence as determined by imaging over time (**Figure 1G**). In HeLa cells with shRNA-mediated depletion of SAMHD1 (**Figure S2A**), the degradation rate of the RNA probe was much lower as compared to control-shRNA treated cells, indicating that RNA degradation in cells is reduced in the absence of SAMHD1 (**Figures 1H and 1I**).

### SAMHD1 depletion causes cellular accumulation of RNA

To determine whether the accumulation of cytosolic RNA was directly caused by the lack of active SAMHD1 in AGS patient fibroblasts, we conducted a rescue experiment. AGS1 cells were transiently transfected with GFP-SAMHD1-WT or RNase-deficient GFP-SAMHD1-Q548X. RNA was quantified using the RNA-specific dye F22, which has excitation and emission properties similar to mCherry, enabling us to measure RNA concentration in cells expressing the GFP-tagged proteins (Li et al., 2006). AGS1 cells overexpressing wild type SAMHD1 showed a decreased amount of cytosolic RNA compared to control-transfected cells (**Figure 2A and 2B**). By contrast, overexpression of mutant SAMHD1 (Q548X) resulted in increased RNA levels compared to cells overexpressing wild type SAMHD1 (**Figures 2A and 2B**). Similar findings were made with transfected HeLa cells or wild type fibroblasts, where shRNA-mediated knockdown of SAMHD1 or overexpression of SAMHD1-Q548X mutant resulted in increased cellular accrual of RNA (**Figures 2C and 2D; Figures S2B and S2C**). HeLa cells expressing control or SAMHD1-shRNA were detected by the expression of a mCherry reporter protein and hence a chemically similar, but green fluorescent RNA-specific dye, E36, was used to quantify RNA instead of F22 (Li et al., 2006). Combined, these findings indicate that RNA accumulation in AGS cells is a direct result of SAMHD1 deficiency.

**Figure 2.**
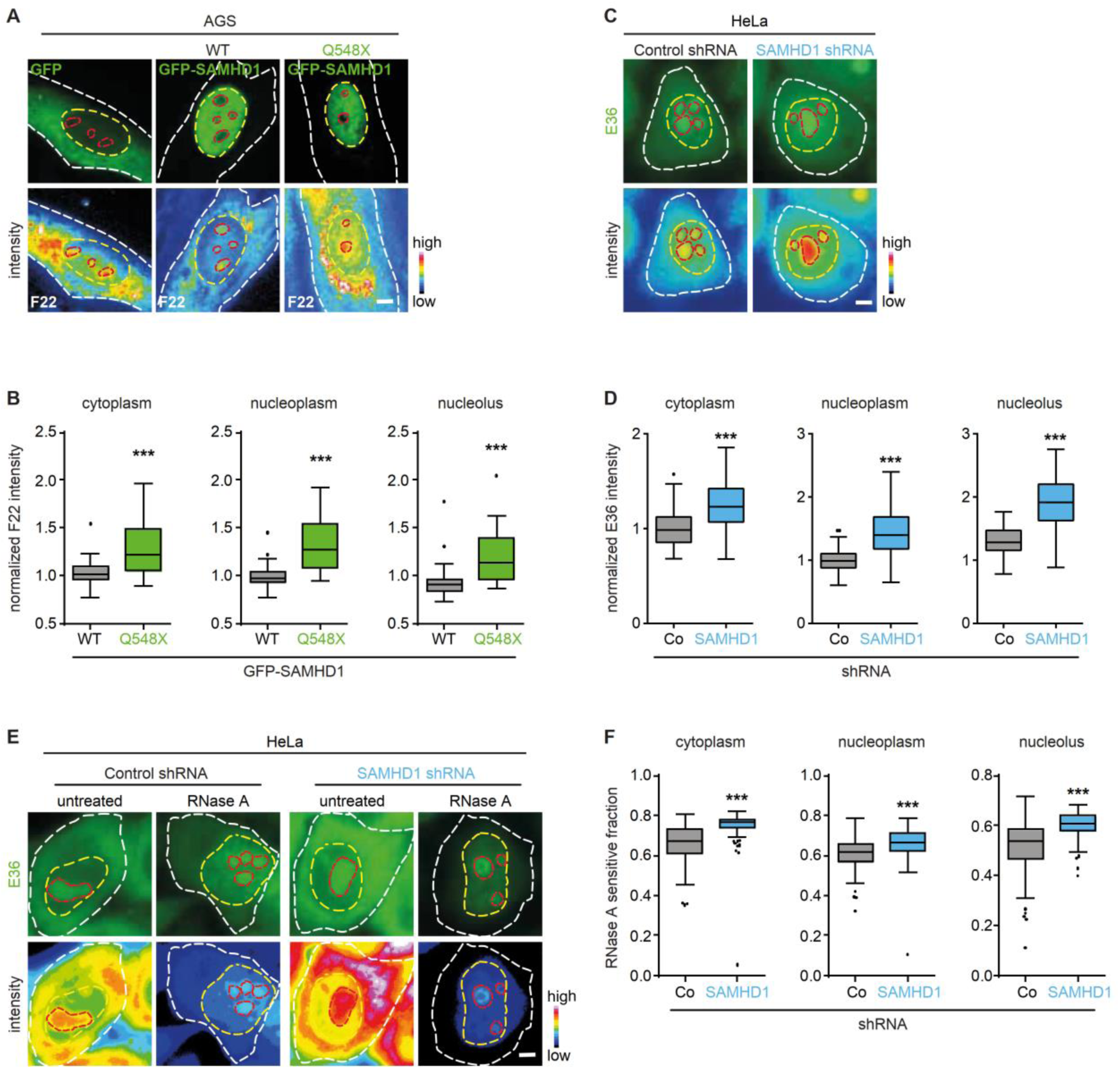
SAMHD1 depletion causes cellular accumulation of RNA. (**A**) Representative images of AGS fibroblasts expressing GFP, GFP-SAMHD1-WT, or GFP-SAMHD1-Q548X (green) and color-coded RNA intensity stained with F22 dye. Outlines of cell, nucleus, and nucleolus are marked with white, yellow and red dashed lines, respectively. Scale bar, 5 µm. (**B**) Quantification of RNA intensity in AGS cells expressing WT or mutant (Q548X) GFP-SAMHD1. n = 46-64 cells. (**C**) Representative images of RNA stained with E36 dye in HeLa cells transfected with control-shRNA or SAMHD1-shRNA along with color-coded intensity images. Outlines of cell, nucleus and nucleolus are marked with white, yellow, and red dashed lines, respectively. Scale bar, 5 μm. (**D**) Quantification of RNA intensity in HeLa cells transfected with control-shRNA or SAMHD1-shRNA. n = 54-58 cells. (**E**) Representative images of RNA stained with E36 dye in HeLa cells transfected with control-shRNA or SAMHD1-shRNA, along with color-coded intensity images, before or after RNase A treatment to deplete accessible ssRNA. Outlines of cell, nucleus, and nucleolus are marked with white, yellow, and red dashed lines, respectively. Scale bar, 5 µm. (**F**) Quantification of RNase A-sensitive ssRNA fraction in HeLa cells transfected with control-shRNA or SAMHD1-shRNA after normalizing the total RNA before RNase A treatment to 1 for each condition. n = 148-203 cells. (**B**, **D**, **F**) Data are representative of at least three independent experiments. Box plots: center line, median; box, interquartile range; whiskers, 1.5x interquartile range; dots, outliers. Mann-Whitney U test. ***, p<0.001.

Given that SAMHD1 degrades ssRNA, we next examined whether SAMHD1-deficient cells specifically accumulate ssRNA. To this end, we stained HeLa cells, transiently expressing SAMHD1-shRNA, with the green fluorescent RNA dye E36, fixed the cells, and subsequently exposed cells to RNase A, which specifically degrades ssRNA. We then determined the E36-positive RNA fraction before and after RNase A treatment. In cells depleted of SAMHD1, a larger proportion of the RNA was accessible to RNase A degradation, when compared to control cells (**Figure 2E**). The relative fraction of soluble ssRNA was 2.3-8% higher in the cytoplasm and 2-11% higher in the nucleus in cells with SAMHD1 depletion compared to control-shRNA treated cells (**Figure 2F**). This led us to conclude that SAMHD1 deficient cells contain higher amounts of ssRNA.

To search for potential binding motifs in SAMHD1 target RNA, we employed photoactivatable ribonucleoside-enhanced crosslinking and immunoprecipitation (PAR-CLIP) (Hafner et al., 2010). HEK293 cells expressing wild type or mutant GFP-SAMHD1 (Q548X and H167Y) were metabolically labelled with the photoactivatable nucleoside 4-thiouridine (4SU) and exposed to UV light to crosslink RNA with nearby RNA-binding proteins. RNA targets crosslinked to SAMHD1 were isolated by anti-SAMHD1 immunoprecipitation and sequenced to identify binding sites on target RNAs marked by nucleotide conversion (Hafner et al., 2010). While very few RNA binding clusters were recovered with wild type SAMHD1, over 8,000 clusters were identified for each mutant most of which mapped to introns (**Figures S3A and S3B**), consistent with the nuclear localization of SAMHD1. However, there was no evidence for preferred binding sites, suggesting that SAMHD1 does not target a specific RNA sequence.

Taken together, our data demonstrate that SAMHD1 has RNase activity both, *in vitro* and *in cellulo*, and that AGS-associated SAMHD1 mutations result in loss of RNase activity, reduced RNA turnover, and cellular accumulation of ssRNA.

### RNA accumulation due to SAMHD1 deficiency perturbs condensates

SAMHD1 deficiency leads to a substantial increase in cellular RNA levels, raising questions about the physiological effects of higher RNA concentration in cells. We had shown in previous work that the overall RNA concentration in cells determines the phase separation behaviour of RNA-binding proteins which are key components of condensates such as stress granules or nucleoli (Maharana et al., 2018). High RNA/protein ratios keep RNA-binding proteins soluble, preventing phase separation, while low RNA/protein ratios promote phase separation into condensates (Banerjee et al., 2017; Maharana et al., 2018; Saha et al., 2016). Further, specific scaffold RNA can compete with non-specific RNA to promote phase separation of RNA-binding proteins and assembly into condensates (Berry et al., 2015; Clemson et al., 2009; Maharana et al., 2018). Given the essential role of RNA in condensate formation, we tested whether the perturbed RNA homeostasis of SAMHD1-deficient cells affects RNA-containing condensates. Immunofluorescence detection of organelle-specific markers in fibroblasts from patient AGS1 and AGS2 (homozygous for H167Y) revealed a marked reduction in arsenite-induced stress granules (G3BP1), nuclear speckles (SC35), and PML bodies (PML) as compared to wild type cells (**Figures 3A-3C**), despite similar expression of the respective constituent proteins (**Figure S4A**). Membraneless compartments were also smaller in size with a less defined morphology (**Figures 3A-3C**), while the condensate-forming proteins were more diffusely distributed in the cytosol, as shown by a reduced enrichment of the nucleolar protein DDX21, stress granule protein G3BP1, and nuclear speckle protein SC35, respectively (**Figure 3D; Figures S4B and S4C**), suggesting impaired condensate assembly in SAMHD1-deficient AGS patient cells.

**Figure 3.**
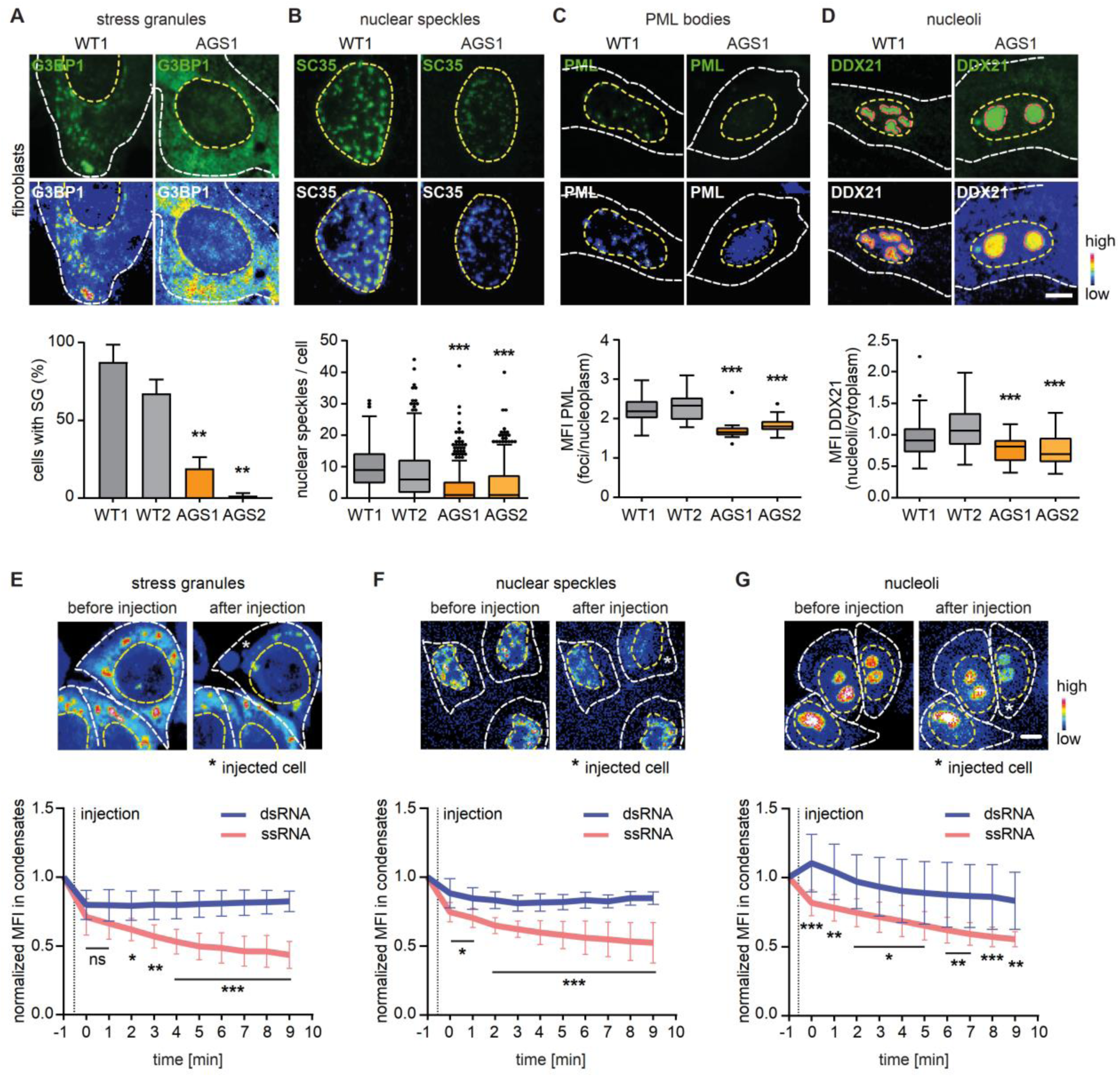
RNA accumulation due to SAMHD1 deficiency perturbs condensates. (**A**-**D**) Stress granules, nuclear speckles, PML bodies, and nucleoli in wild type (WT1) and AGS1 fibroblasts stained with indicated marker proteins (green) along with color-coded intensity. Outlines of cell and nucleus are marked with white and yellow dashed lines, respectively. Scale bars, 5 μm. Quantification of membraneless compartment perturbation in wild type (WT1, WT2) and AGS (AGS1, AGS2) fibroblasts. G3BP1 shown as percent of cells with stress granules, n = 94-135 cells; SC35, n = 222-559 cells; PML, n = 30-95 cells; DDX21, n = 59-125 cells. Box plots: center line, median; box, interquartile range; whiskers, 1.5x interquartile range; dots, outliers. Kruskal-Wallis test with Dunn’s multiple comparisons test. ***, p<0.001. Data are representative of three independent experiments. (**E**-**G**) Representative images of HeLa cells expressing G3BP1-mCherry (stress granules), SRRM1-GFP (nuclear speckles), or nucleolin-GFP (nucleoli) before and after microinjection of a 45 nt ssRNA or a 45 bp dsRNA oligonucleotide. Microinjected cells are marked with stars. Outlines of cell and nucleus are marked with white and yellow dashed lines, respectively. Scale bars, 5 μm. Quantification of mean fluorescence intensities (MFI) of marker proteins (G3BP1, SRRM1, nucleolin) within stress granules, nuclear speckles and nucleoli after microinjection of ssRNA or dsRNA. MFI of marker proteins was normalized to 1 for the time point just prior to microinjection and corrected for bleaching by using the non-injected cell in its vicinity. n = 6-11 cells. Mean ± SD. Two-way ANOVA with Sidak’s multiple comparisons test. *,p<0.05; **, p<0.01; ***, p<0.001. (**A-G**) Representative data from at least two independent experiments.

Since AGS cells are chronically depleted of SAMHD1, we next sought to determine whether the observed condensate abnormalities were directly dependent on SAMHD1. Like in AGS cells, in HeLa cells in which SAMHD1 was acutely downregulated by shRNA expression, formation of stress granules was impaired (**Figure S4D**). Again, nuclear speckles and nucleoli contained lower amounts of their constituent proteins as shown by staining of SC35 and fibrillarin (FIB) or DDX21 (**Figures S4E-S4G**), respectively, consistent with abnormal phase separation and confirming that RNA accumulation due to SAMHD1 deficiency disturbs formation of cellular condensates.

To test if the observed changes in condensate formation compromise condensate function, we focussed on post-transcriptional modifications of RNA, which take place in nucleoli and speckles (Boulon et al., 2010; Galganski et al., 2017). Mass spectrometry of nucleoside digests obtained from total RNA of HeLa cells with siRNA-mediated knockdown of SAMHD1 (Kretschmer et al., 2015), in which nucleoli and speckles are perturbed, revealed profound changes in methylation, acetylation, and pseudouridinylation (**Figure S5**). Similarly, SAMHD1-deficient patient cells showed an increased overall 2’-O-methylation of rRNA (**Figure S6**), as assessed by RiboMethSeq analysis (Marchand et al., 2016), confirming that loss of proper formation of nucleoli and nuclear speckles due to SAMHD1 deficiency causes global alterations in post-transcriptional RNA modifications that could affect the innate immune system. Altogether, these results establish that increased cellular RNA concentrations due to a lack of SAMHD1 impede normal formation of RNA condensates and cause mislocalization of RNA-binding proteins.

To assess whether global condensate perturbation was specific for SAMDH1-deficient cells, we analyzed condensate behavior in primary fibroblasts from AGS patients carrying mutations in TREX1, encoding a DNase, and RNASEH2 (**Fig. S7A-C**). PML body formation was not altered in patient cells compared to two wild type controls (**Fig. S7B**). Stress granule formation in TREX1-deficient cells was reduced by 9%, while recruitment of fibrillarin into nucleoli was reduced by 11% in RNASEH2-deficient cells. However, we did not observe a global breakdown of membraneless compartments in AGS fibroblasts harboring TREX1 or RNASEH2 mutations (**Fig. S7A-C**) as seen in SAMHD1-deficient cells. We also assessed condensate formation in HeLa cells with knockdown of the following cellular RNases active in different cellular compartments (**Fig. S7D**): XRN1, the major cytosolic 5’-3’ exoribonuclease involved in mRNA degradation and processing of rRNAs (Blasco-Moreno et al., 2019); EXOSC10, part of the nuclear exosome involved in accurate maturation of stable RNA species such as rRNA and in the elimination of antisense RNA species or defective mRNAs (van Dijk et al., 2007); RNASET2, which cleaves microbial and mitochondrial RNA in endolysosomes (Greulich et al., 2019). Overall, we did not observe a global disturbance of condensate formation in cells with depletion of XRN1, EXOSC10, or RNASET2, as seen in SAMHD1-deficient cells. While knockdown of XRN1 had no effect on stress granule formation, knockdown of EXOSC10 and RNASET2 was associated with reduced enrichment of G3BP1 in stress granules (**Fig. S7E**), that was not as pronounced as in SAMHD1-depleted cells (**Fig. S4B**). Thus, SAMHD1 depletion led to approximately 45% downregulation of G3BP1 enrichment whereas in case of EXOSC10, RNASET2, and XRN1, the reduction of G3BP1 enrichment was only 15%, 17%, and 3%, respectively. PML bodies were not affected by knockdown of XRN1 or EXOSC10, although PML enrichment was slightly reduced by 6% in cells depleted of RNASET2 (**Fig. S7F**). In contrast, depletion of EXOSC10 or RNASET2 had no effect on fibrillarin enrichment in nucleoli, while knockdown of XRN1 was associated with a 20% reduced enrichment of fibrillarin in nucleoli (**Fig. S7G**). These findings may be due to changes in cellular RNA concentrations, or may reflect stress responses, as both, PML bodies and nucleoli are sensitive to RNA stress (Boulon et al., 2010). Notwithstanding, these data suggest that global impairment of condensate formation is specifically seen only in SAMHD1-deficient cells.

### ssRNA, but not dsRNA, dissolves condensates

To further test whether an acute increase in cellular ssRNA levels is the underlying cause of the observed condensate abnormalities, we increased the cellular concentration of ssRNA - the substrate of SAMHD1 - by microinjection of a 45 nt ssRNA oligonucleotide into HeLa cells constitutively expressing mCherry-tagged stress granule protein G3BP1, GFP-tagged nuclear speckle protein SRRM1, or GFP-tagged nucleolar protein nucleolin (Poser et al., 2008). The integrity of stress granules, nuclear speckles and nucleoli was impaired as seen by the loss of condensate-residing proteins, with only 50% of marker protein retained in condensates within 10 min after microinjection (**Figures 3E-3G**). In agreement with this, previous work showed that stress granules can be dissolved by microinjection of tRNA into the cytoplasm (Maharana et al., 2018). We next investigated the role of RNA strandedness on condensate dissolution by comparing the effects of either ssRNA or dsRNA. Microinjection of ssRNA caused a decrease in the amount of constituent proteins inside stress granules, nuclear speckles, and nucleoli, while microinjection of dsRNA had only little effect on condensate integrity (**Figures 3E-3G**).

Given the increased relative RNase-sensitive fraction of soluble ssRNA in SAMHD1-depleted cell of 2.3-8% in the cytoplasm and 2-11% in the nucleus compared to control-shRNA treated cells (**Fig. 2F**), we sought to assess whether the amount of microinjected RNA corresponds to the observed increase in cellular ssRNA. We therefore estimated the amounts RNA used in our microinjection experiments in HeLa cells (**Fig. S8**). On average, 4.6 fl were injected with a range of 0.6 -17.8 fl, depending on the inner capillary diameter which is 0.5 µm ± 0.2 µm according to the manufacturer. This corresponds to an average RNA amount of 0.0055 pg, reflecting ∼0.06%, and ∼0.5% of cytoplasmic and nuclear RNA levels, respectively. Thus, the amounts of RNA used in our microinjection experiments are within the physiological range of the observed ssRNA increase in SAMHD1-deficient cells and support the notion that an acute increase of even small amounts of cellular ssRNA is sufficient to impair condensate formation.

To validate our *in cellulo* observations further, we examined the effect of ssRNA and dsRNA on RNA condensate assembly *in vitro*. We reconstituted stress granule-like condensates in HeLa cell extract by adding recombinant G3BP1, an important scaffold protein for stress granule assembly in cells (Freibaum et al., 2021; Guillén-Boixet et al., 2020). We then added fluorescently labelled ssRNA or dsRNA to quantify the effect of RNA strandedness on phase behaviour. At lower concentrations, both ssRNA and dsRNA were enriched within condensates (**Figures 4A and 4B**). However, at a concentration of 5 μM of ssRNA, condensates dissolved with only ∼12% of initial G3BP1 remaining in condensates, while the same concentration of dsRNA had little effect on condensate integrity with ∼75% of initial G3BP1 still contained in stress granule-like condensates (**Figures 4A and 4B**).

**Figure 4.**
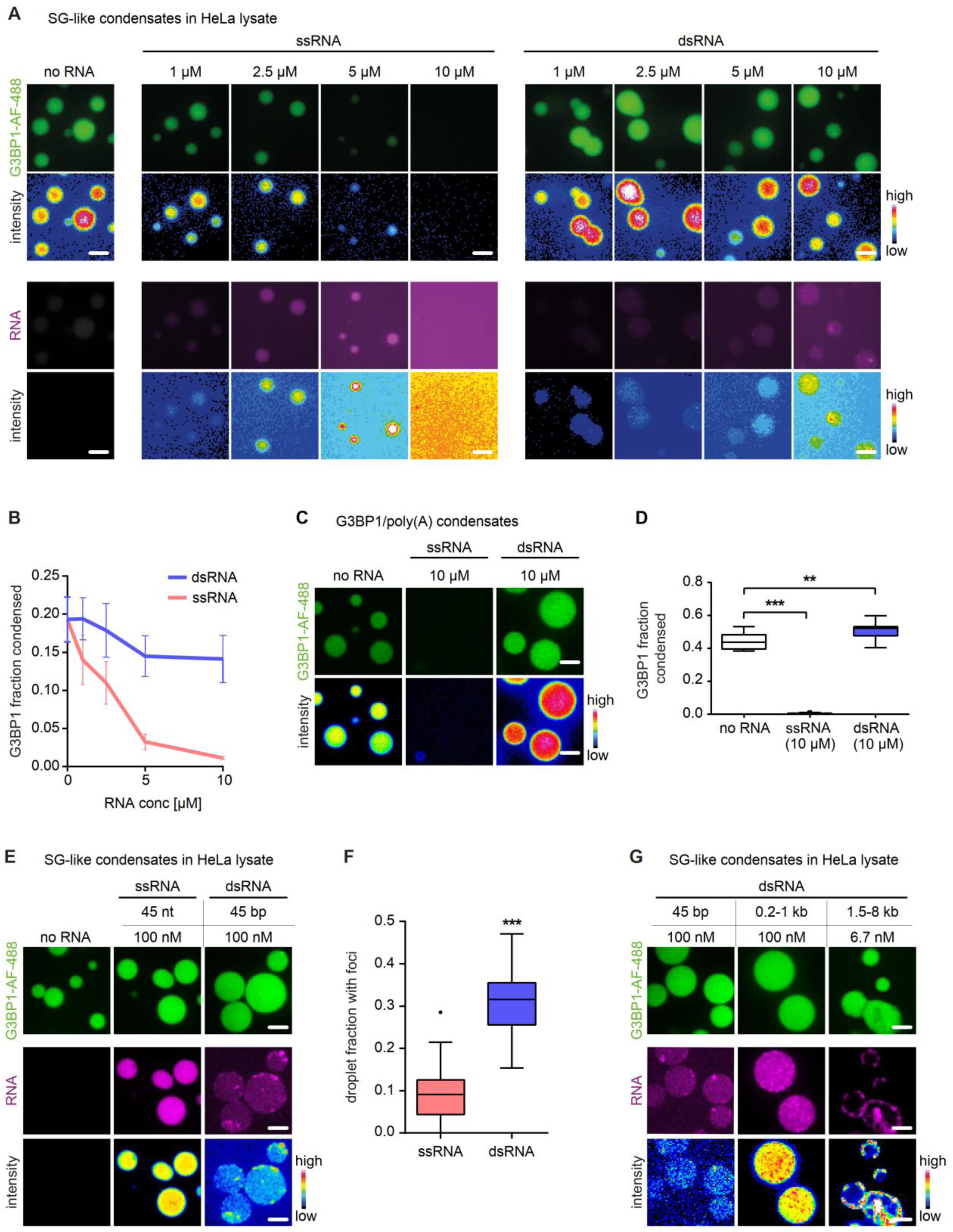
ssRNA, but not dsRNA, dissolves *in vitro* reconstituted stress granule-like condensates. (**A**) Representative images of stress granule-like condensates induced by addition of AF-488-labeled SNAP-G3BP1 to HeLa cell lysates to which either Cy5-labeled ssRNA or dsRNA was added, along with color-coded intensity images of RNA and G3BP1 in stress granule-like condensates. Scale bars, 5 µm. (**B**) Quantification of G3BP1 fraction in stress granule-like condensates (cumulative G3BP1 intensity in condensates per total G3BP1). n =1000-1450 condensates. Mean ± SD. (**C**) Representative images of minimal G3BP1 condensates induced by adding poly(A)-RNA to AF 488-labeled SNAP-G3BP1 in physiological buffer to which ssRNA or dsRNA was added. Scale bars, 5 µm. (**D**) Quantification of G3BP1 fraction within G3BP1 condensates in presence or absence of ssRNA or dsRNA. n = 30-45 field of views. Box plots: center line, median; box, interquartile range; whiskers, 1.5x interquartile range; dots, outliers. Kruskal-Wallis test with Dunn’s multiple comparisons test. **, p<0.01; ***, p<0.001. (**E**) Representative images of stress granule-like condensates induced by addition of AF 488-labeled SNAP-G3BP1 to HeLa cell lysates to which either Cy5-labeled ssRNA (45 nt) or dsRNA (45 bp) was added, along with color-coded intensity images of G3BP1. Scale bars, 5 µm. (**F**) Quantification of proportion of stress granule-like condensates containing RNA aggregates. n = 25 field of views. Box plots: center line, median; box, interquartile range; whiskers, 1.5x interquartile range; dots, outliers. Mann-Whitney U test. ***, p<0.001. (**G**) Representative images of aggregates in stress granule-like condensates build after mixing AF 488-labeled SNAP-G3BP1 and dsRNA of increasing size (Cy5-labeled 45 bp dsRNA, Rhodamine-labeled low molecular weight or high molecular weight poly(I:C)) in HeLa cell lysate along with color-coded RNA intensity. Scale bars, 5 µm.

Because stress granules formed in HeLa cell extract contain multiple RNA-binding proteins and G3BP1 has been suggested to bind both ssRNA and dsRNA (Kim et al., 2019; Protter and Parker, 2016), we next sought to determine whether RNA also affects minimal stress granules formed from only two components, the scaffold protein G3BP1 and RNA. To this end, we reconstituted minimal stress granules *in vitro* by mixing single-stranded poly(A)-RNA and recombinant G3BP1 in a physiological buffer (Guillén-Boixet et al., 2020) and then added 10 μM of ssRNA or dsRNA (**Figure 4C**). Consistent with our findings with HeLa cell extract, ssRNA prevented formation of G3BP1 condensates, whereas dsRNA enhanced G3BP1 phase separation by another 10% (**Figures 4C and 4D**). Thus, ssRNA but not dsRNA readily prevents assembly of stress granule-like condensates, presumably by impairing G3BP1-mediated interactions that are required for stress granule formation.

Interestingly, while ssRNA was uniformly enriched in stress granule-like condensates at low concentrations, dsRNA assembled into non-homogenous assemblies or foci in a large fraction of condensates (**Figures 4A, 4E-4F**), which may help in sequestering dsRNA more efficiently as larger particles are slower to diffuse. However, dsRNA of similar length did not form any foci in G3BP1/poly(A) condensates (**Figure 4C**) suggesting that dsRNA foci formation in stress granule-like condensates does not only involve intermolecular interactions between dsRNA molecules as reported previously for other RNAs (Matheny et al., 2021), but requires additional RNA-binding proteins present in cell lysates. As stress granule-like condensates are equally complex as native stress granules and contain a large number of different RNA-binding proteins (Freibaum et al., 2021), we speculate that yet unknown RNA-binding proteins can enrich and sequester dsRNA inside stress granules. The size of these dsRNA assemblies increased with the size of the dsRNA (**Figure 4G**), suggesting also a role of RNA as a scaffold. Collectively, these findings reveal opposing effects of ssRNA and dsRNA on RNA-binding protein-mediated phase separation. While ssRNA promotes dissolution of condensates, dsRNA is sequestered inside condensates in large assemblies.

### Cellular condensates sequester dsRNA

dsRNA acts as major inducer of cytosolic RNA sensors engagement of which triggers type I IFN signalling, a hallmark of AGS. Our *in vitro* experiments demonstrated that ssRNA dissolves condensates, while dsRNA becomes enriched in condensates. This led us to hypothesize that cellular condensates may serve an important immunoregulatory function by sequestering endogenous dsRNA. Given impaired condensate formation in AGS patient fibroblasts and in HeLa cells with SAMHD1 depletion, we investigated the subcellular distribution of endogenous dsRNA in condensates *in cellulo*. We immunostained dsRNA using the J2 antibody (Dhir et al., 2018) in HeLa cells that were stressed with sodium arsenite to induce stress granule formation. Immunostaining with J2 showed that a large fraction of the cytoplasmic signal colocalized with mitochondria (**Figure S9**), in agreement with mitochondrial transcription being a major source of cellular dsRNA (Dhir et al., 2018). Because the relevant immunosensors for dsRNA reside in the cytosol, we excluded the mitochondrial J2 signal from image analysis (**Figure S9**). Performing this analysis and using eIF3η as a marker for stress granules, we found that significant amounts of dsRNA were enriched in stress granules (**Figures 5A-5B**), consistent with the notion that stress granules can sequester large amounts of dsRNAs. The origin of these dsRNAs is not clear, but previous work has shown that many nuclear RNAs leak into the cytoplasm during stress (Holley et al., 2015; Zander et al., 2016). We also analysed HeLa cells for dsRNA enrichment in speckles using SSRM1-GFP as a marker and found that nuclear speckles also sequester dsRNA (**Figures 5C-5D**).

**Figure 5.**
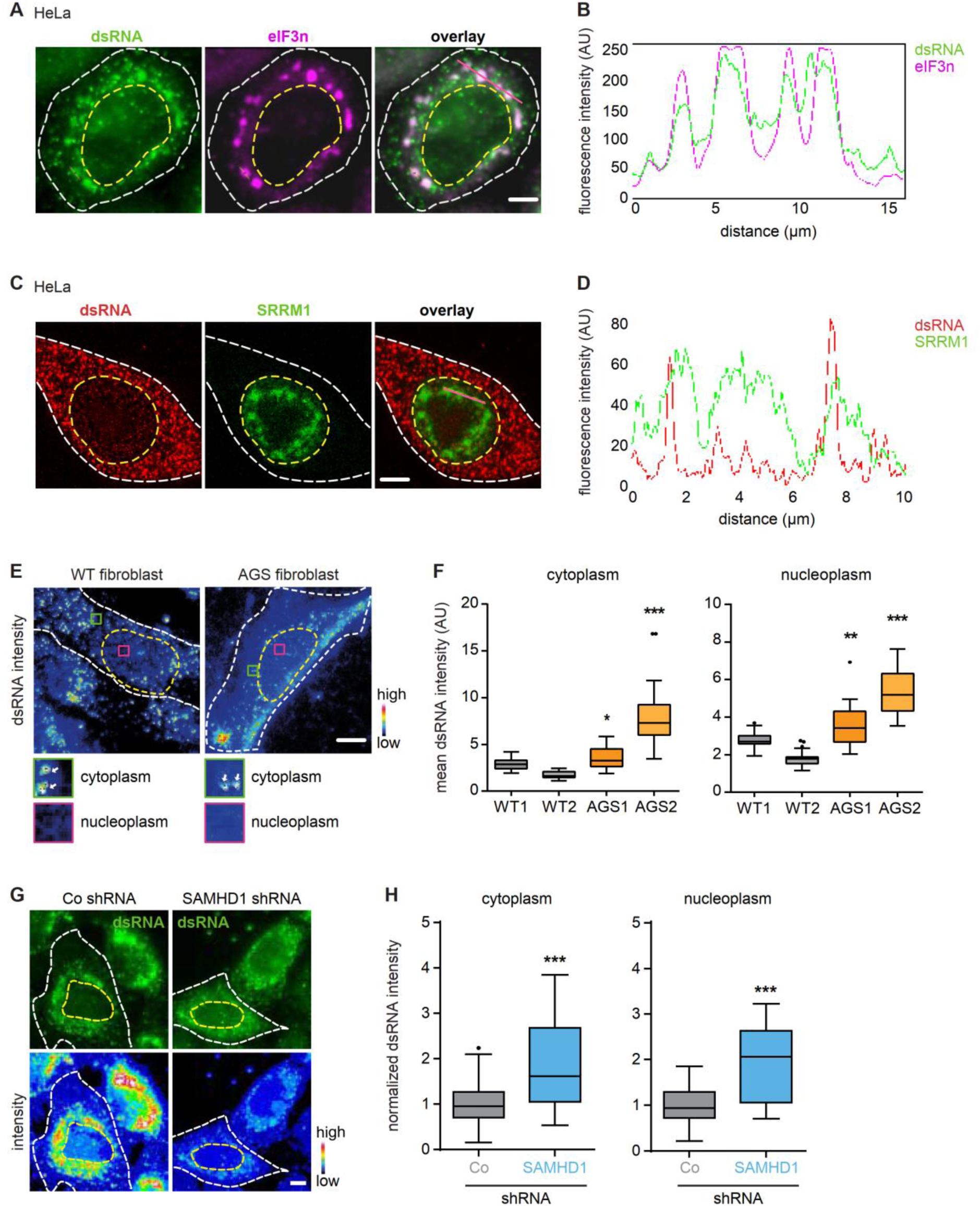
Stress granules sequester self dsRNA. (**A**, **B**) Representative images of arsenite-stressed HeLa cells stained with the stress granule marker eiF3η, dsRNA stained with J2. White dotted lines mark the cellular boundary and yellow dotted lines denote the nuclear outline. Scale bar, 5 µm. Colocalization of dsRNA and stress granules as shown by fluorescence density profiles of dsRNA and eiF3η signals corresponding to region marked with pink line in (**A**). (**C**, **D**) Representative images of HeLa cells expressing the nuclear speckle marker SRRM1-GFP, dsRNA stained with J2. White dotted lines mark the cellular boundary and yellow dotted lines denote the nuclear outline. Scale bar, 5 µm. Colocalization of dsRNA and nuclear speckles as shown by fluorescence density profiles of dsRNA and SRRM1 signals corresponding to region marked with pink line in (**C**). (**E**) Representative color-coded intensity images of AGS and WT cells showing J2-stained dsRNA. White arrows in insets show mitochondrial dsRNA staining, which was omitted during quantification. White dotted lines mark the cellular boundary and yellow dotted lines denote the nuclear outline. Scale bar, 5 µm. (**F**) Quantification of J2-stained dsRNA in wild type (WT1, WT2) and patient cells (AGS1, AGS2). n = 19-37 cells. Box plots: center line, median; box, interquartile range; whiskers, 1.5x interquartile range; dots, outliers. Kruskal-Wallis test with Dunn’s multiple comparisons test. *, p<0.05; **, p<0.01; ***, p<0.001. (**G**) Representative images of mean fluorescence intensity of dsRNA in HeLa cells expressing control-shRNA or shRNA against SAMHD1 along with color-coded intensity. Outlines of cells and nuclei are marked with white and yellow dashed lines, respectively. Scale bar, 5 μm. (**H**) Quantification of mean fluorescence intensity of dsRNA in SAMHD1-depleted HeLa cells normalized to control-shRNA. n = 118-131 cells. Box plots: center line, median; box, interquartile range; whiskers, 1.5x interquartile range; dots, outliers. Mann-Whitney U test. ***, p<0.001. (**A-H**) Data are representative of three independent experiments.

Our data above showed that loss of SAMHD1 in AGS patient cells causes accumulation of ssRNA and impairs the assembly of condensates, including stress granules and nuclear speckles. We thus predicted that compromised condensate assembly leads to higher cytosolic dsRNA levels in cells lacking SAMHD1. To examine this further, we performed immunostaining of dsRNA in SAMHD1-deficient AGS cells and in HeLa cells with SAMHD1 depletion both of which revealed a higher J2 signal in the nucleoplasm and cytosol compared to controls (**Figures 5E-5H**), confirming that endogenous dsRNA levels are increased in SAMHD1-deficient cells.

Considering that SAMHD1 is specific for ssRNA, the accumulation of dsRNA in SAMHD1-deficient cells is surprising and requires an explanation. We hypothesized that the high amount of cytosolic dsRNA in SAMHD1-deficient cells could be due to a release of dsRNA from disintegrating condensates. To test the idea that endogenous dsRNA can be released from aberrantly formed condensates, we overexpressed the mitotic kinase DYRK3, a condensate dissolvase that was previously shown to dissolve nuclear splicing speckles as well as cytosolic stress granules (Rai et al., 2018). In line with the solubilising effect of DYRK3 on nuclear speckles, DYRK3 expression increased dsRNA levels in the nucleus (**Figure 6A and 6B**). In the absence of DYRK3, cytosolic dsRNA was enriched in stress granules (**Figures 6C and 6D**). However, in DYRK3-expressing cells, stress granule formation was impaired and this was accompanied by an increase of dsRNA outside of stress granules by up to 50% as compared to cells that did not overexpress DYRK3 (**Figures 6C and 6D**). Similarly, cytosolic dsRNA levels were increased in HeLa cells in which stress granule formation was inhibited specifically by treatment with the integrated stress response inhibitor (ISRIB) (Sidrauski et al., 2015) (**Figures 6E and 6F**). Thus, lack of dsRNA sequestration in membraneless compartments causes cytosolic accrual of dsRNA.

**Figure 6.**
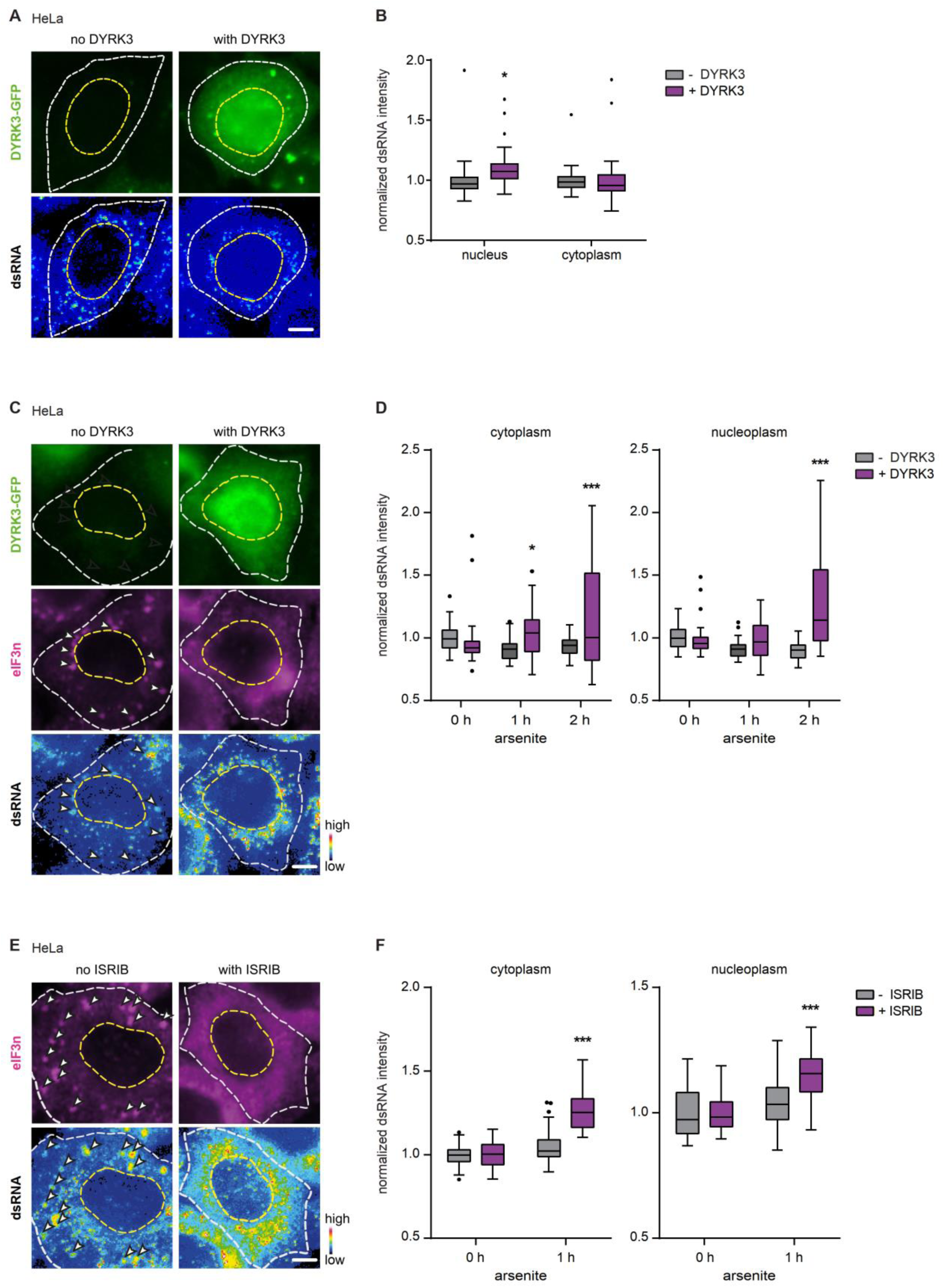
Dissolution of condensates causes cytosolic accrual of dsRNA. (**A**) Representative images of HeLa cells, expressing DYRK3-GFP, stained with anti-dsRNA J2 antibody. Outlines of cell and nucleus are marked with white and yellow dashed lines, respectively. Scale bar, 5 µm. (**B**) Quantification of dsRNA in nucleus and cytoplasm of DYRK3-GFP expressing HeLa cells. Fluorescence intensities were normalized to no DYRK3 expression. Box plots: center line, median; box, interquartile range; whiskers, 1.5x interquartile range; dots, outliers. n = 32-46 cells. Two-way ANOVA with Sidak’s multiple comparisons test. *, p<0.05. (**C**) Representative images of arsenite-stressed HeLa cells transiently expressing or not expressing DYRK3-GFP immunostained with stress granule marker eIF3η along with color-coded intensity images of dsRNA stained with J2 antibody. The white arrow heads mark stress granules in cytoplasm. Scale bar, 5 µm. (**D**) Quantification of dsRNA intensity in HeLa cells after 1 or 2 h of arsenite stress, expressing or not expressing DYRK3-GFP. The dsRNA intensity measured before application of stress was normalized to 1. n= 24-78 cells. Box plots: center line, median; box, interquartile range; whiskers, 1.5x interquartile range; dots, outliers. One-way ANOVA with Sidak’s multiple comparisons test. *, p<0.05; ***, p<0.001. (**E**) Representative images of arsenite-stressed HeLa cells treated with stress granule formation inhibitor ISRIB or empty vehicle and immunostained for stress granule marker eIF3η. dsRNA stained using J2 antibody (intensity color-coded). White dotted lines mark the cellular boundary and yellow dotted lines denote the nuclear outline. The white arrow heads mark stress granules in cytoplasm. Scale bar, 5 µm. (**F**) Quantification of dsRNA intensity in HeLa cells after 1 h of arsenite stress, with or without ISRIB treatment. The dsRNA intensity measured before treatment was normalized to 1. n = 30-49 cells. Box plots: center line, median; box, interquartile range; whiskers, 1.5x interquartile range; dots, outliers. Two-way ANOVA with Sidak’s multiple comparisons test. ***, p<0.001. (**A**-**F**) Data are representative of at least two independent experiments.

### RNA sensing pathways are activated in SAMHD1 deficiency

A predominant feature of the neuroinflammatory disease AGS is constitutive activation of type I IFN signalling triggered by aberrant immune sensing of self nucleic acids (Lee-Kirsch, 2017; Schlee and Hartmann, 2016). Although type I IFN activation in SAMHD1 deficiency has been attributed to cGAS-dependent immune recognition of dsDNA (Coquel et al., 2018; Maelfait et al., 2016), we hypothesized that activation of RNA sensing pathways due to RNA accrual could also account for type I IFN activation in AGS. To dissect the nucleic acid sensing pathways mediating type I IFN activation in SAMHD1-deficient cells, we measured the expression of IFN-stimulated genes in AGS1 fibroblasts treated with siRNA targeting either the dsDNA sensor cGAS or the dsRNA sensors MDA5 and RIG-I, respectively. Knock down of either cGAS, MDA5 or RIG-I diminished type I IFN signalling as shown by reduced expression of the IFN-induced protein MX1 and IFN-stimulated genes (**Figure 7A and 7B**), suggesting that type I IFN signalling in SAMHD1 deficiency is not only triggered by dsDNA (Coquel et al., 2018; Maelfait et al., 2016), but also by dsRNA. We validated this further in HeLa cells with CRISPR/Cas9-mediated knockout of MDA5, RIG-I or both. Cells were treated with control or SAMHD1-siRNA and then stimulated with RIG-I ligand (5’ppp dsRNA) or MDA5 ligand obtained from encephalomyocarditis virus-infected Vero cells (EMCV) (**Figure 7C and 7D**). The extent of type I IFN signalling was measured by determining IP-10 (CXCL10) secretion. In response to the RIG-I ligand 5’ppp dsRNA, IP-10 secretion was increased in MDA5^-/-^ cells treated with SAMHD1-siRNA compared to cells treated with control-siRNA, but secretion was completely abrogated in RIG-I^-/-^ and RIG-I^-/-^/MDA5^-/-^ double knockout cells (**Figure 7C**), consistent with enhanced RIG-I-mediated signalling in SAMHD1-depleted cells. On the other hand, when cells were stimulated with the MDA5 ligand EMCV, IP-10 secretion was only partially decreased in MDA5^-/-^ cells, but completely blunted in RIG-I^-/-^/MDA5^-/-^ double knockout cells (**Figure 7D**), suggesting enhanced RIG-I activation by dsRNA in the absence of MDA5 (Kato et al., 2008) due to SAMHD1 knockdown. These findings are in line with the notion that both RIG-I and MDA5 contribute to type I IFN activation in SAMHD1 deficiency.

**Figure 7.**
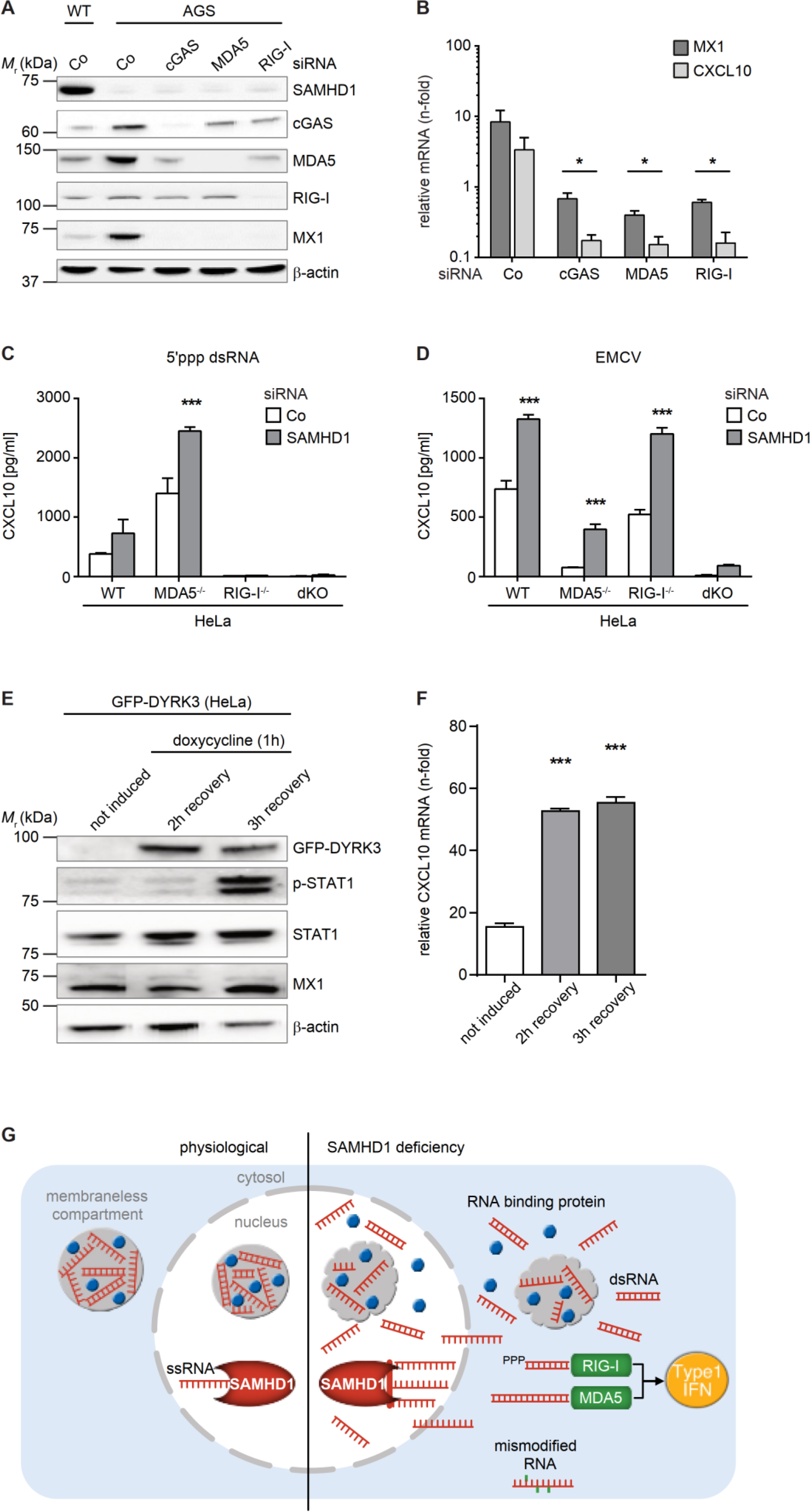
RNA sensing pathways are activated in SAMHD1 deficiency. (**A**) Western blot of wild type (WT) and AGS fibroblasts treated with control-siRNA (Co) or siRNA against cGAS, MDA5, or RIG-I, immunostained with the indicated antibodies. (**B**) Expression of IFN-stimulated genes in AGS cells treated with indicated siRNA, relative to wild type cells treated with control-siRNA. Mean ± SEM of three independent experiments run in triplicate technical replicates. One-way ANOVA with Dunnett’s multiple comparisons test, co-siRNA vs. sensor-siRNA. *, p<0.05. (**C**, **D**) IP-10 secretion of wild type HeLa cells (WT) or HeLa cells with knockout of MDA5 (MDA5^-/-^) or RIG-I (RIG-I^-/-^) or with double knockout of MDA5 and RIG-I (MDA5^-/-^/RIG-I^-/-^; dKO) treated with control-siRNA or SAMHD1-siRNA after stimulation with 5’ppp dsRNA (**C**) and EMCV total RNA (**D**). Mean ± SEM of two independent experiments with four technical replicates. Two-way ANOVA with Sidak’s multiple comparisons test. ***, p<0.001. (**E**) Representative Western blot of GFP-DYRK3 expressing HeLa cells without or with 1 h doxycycline treatment showing induction of p-STAT1 and MX1 after 3 h of recovery. (**F**) Expression of CXCL10 mRNA in GFP-DYRK3 expressing HeLa cells either untreated or treated with doxycycline for 1 h, followed by 2 h or 3 h recovery. Mean ± SEM of triplicate technical replicates, representative of three independent experiments. One-way ANOVA with Dunnett’s multiple comparisons test, doxycycline untreated vs. treated, ***, p<0.001. (**G**) Model: in normal cells, the RNase activity of SAMHD1 controls cellular levels of ssRNA, which together with RNA-binding proteins, are required for condensate formation by phase separation. In SAMHD1 deficiency, unmetabolized ssRNA accumulates, impeding formation of condensates which sequester dsRNA. This leads to release of dsRNA from condensates with subsequent activation of RLR-dependent type I IFN signalling.

### Disruption of condensate formation triggers a type I interferon response

Our data indicate that dsRNA accumulation causes constitutive type I IFN activation in SAMHD1 deficient cells. We thus hypothesized that experimentally induced condensate dissolution and subsequent dsRNA release should also activate type I IFN signalling. Indeed, condensate dissolution induced by doxycycline-regulated expression of DYRK3-GFP (Rai et al., 2018) led to induction of type I IFN signalling in HeLa cells as shown by enhanced expression of the IFN-induced protein MX1 and phosphorylation of STAT1 (**Figure 7E**) as well as upregulation of the IFN-stimulated gene CXCL10 (**Figure 7F**). Thus, impaired confinement of immunogenic RNA to condensates triggers aberrant activation of the innate immune system. Collectively, these results reveal an important role for condensates in sequestering immunostimulatory dsRNA and show that loss of condensate integrity with subsequent dsRNA release can cause autoinflammatory disease.

## Discussion

Bona fide recognition of viral RNA through pattern recognition receptors is challenged by the vast number of potentially immunogenic endogenous RNAs in cells. Our understanding as to how cytosolic RNA sensors accurately discriminate between self and nonself RNA to avoid aberrant immune activation has remained incomplete. In this work, we demonstrate that condensates sequester immunostimulatory self RNA, thereby preventing inappropriate immune activation. We show that loss-of-function mutations in the RNase SAMHD1 associated with autoinflammatory disease cause cellular accumulation of RNA and impaired assembly of RNA-containing condensates. This in turn causes the release of immunogenic dsRNA from condensates, activating the innate immune system (**Figure 7G**). Thus, sequestration of endogenous RNA inside condensates is an important mechanism of cellular self RNA tolerance, which leads to autoinflammation when RNA sequestration in condensates is defective.

The question as to whether SAMHD1 functions as RNase is controversially discussed and has remained an unresolved question in the field. While SAMHD1 was reported to function as RNase (Beloglazova et al., 2013; Ryoo et al., 2014), this idea was challenged in other studies (Antonucci et al., 2016; Goldstone et al., 2011; Goncalves et al., 2012; Seamon et al., 2015; Welbourn and Strebel, 2016). Using various physiological and synthetic substrates, we now demonstrate that SAMHD1 acts as a ssRNA 3’-exonuclease capable of degrading a broad range of cellular ssRNA species in a sequence-independent manner. Although mass spectrometric analysis of purified recombinant SAMHD1 did not reveal any contaminating RNase species, we cannot fully rule out the presence of an RNase contaminant. However, AGS-associated SAMHD1 mutants do not exhibit any detectable ribonucleolytic activity, implicating a lack of RNase function of SAMHD1 in disease pathogenesis. In line with our *in vitro* findings, loss of SAMHD1 decreases RNA turnover in AGS patient cells causing cellular accrual of RNA. The ensuing perturbation of global RNA homeostasis leads to cytosolic accumulation of immunostimulatory self RNA by two mutually dependent mechanisms. First, failure to degrade ssRNA due to lack of functional SAMHD1 leads to increased cellular concentrations of ssRNA impeding the formation of condensates such as nucleoli, nuclear speckles, or stress granules. This can be explained by the fact that RNA-protein condensates form by the process of liquid-liquid phase separation, which is very sensitive to changes in RNA concentration (Alberti and Dormann, 2019; Banani et al., 2017; Maharana et al., 2018). Importantly, the local concentration of RNA critically affects the phase behaviour of scaffold RNA-binding proteins and thus their threshold of condensation, also referred to as the saturation concentration (Berry et al., 2015; Henninger et al., 2021; Maharana et al., 2018). Secondly, RNA-protein condensates store immunostimulatory dsRNA, which is released when condensates are compromised leading to RLR-dependent activation of type I IFN signalling (**Figure 7G**).

Interestingly, we demonstrate distinctive effects of RNA on condensate formation depending on the structural properties of the RNA. While high concentrations of ssRNA impede condensation or dissolve existing condensates, dsRNA is sequestered into condensates without significantly affecting condensation. Given that dsRNA, and not ssRNA, acts as ligand for cytosolic RNA sensors such as RIG-I or MDA5, our observation that stress granules become enriched in dsRNA provides a mechanism by which immunostimulatory self RNA could be cleared from the cytosol during stress. In unstressed cells, dsRNA might be preferentially sequestered by steady condensates such as nuclear speckles or nucleoli. However, we did not observe enrichment of dsRNA in the nucleolus, which could be due to the inaccessibility of its dense structure by the J2 antibody. In support of this notion, DYRK3-driven dissolution of stress granules and nuclear speckles led to an increase in dsRNA, both in the cytoplasm and nucleus, supporting a crucial role of cytosolic and nuclear condensates in keeping inappropriate immune responses to self dsRNA in check. As such, the RNase activity of SAMHD1 exerts a critical housekeeping function required to maintain an optimally balanced RNA metabolism thus ensuring self RNA tolerance.

Our results also demonstrate a pivotal role of RNA concentration for organizing the RNA-rich cellular environment of the cytosol and nucleoplasm by condensate formation. While sequestration of endogenous dsRNA into condensates protects the cytosol from erroneous innate immune activation, such a mechanism may also explain how RNA viruses escape immune surveillance by replicating in RNA condensates, as has recently been demonstrated for vesicular stomatitis virus or SARS-CoV-2 (Heinrich et al., 2018; Perdikari et al., 2020; Wang et al., 2021).

The formation of aberrant condensates and protein aggregates has been linked to neurodegenerative diseases such as amyotrophic lateral sclerosis (ALS) and frontotemporal dementia (FTD) (Molliex et al., 2015; Patel et al., 2015). One protein that is frequently found in an abnormally aggregated state in these disorders is the nuclear protein TDP-43 (Mackenzie and Rademakers, 2008). Interestingly, TDP-43-deficient cells were recently shown to accumulate dsRNA leading to type I IFN activation and cell death (Dunker et al., 2021). This suggests that specific RNA-binding proteins such as TDP-43 could function as RNA-sequestering molecules that confine dsRNA to specific condensates. Importantly, our findings demonstrate that not only enhanced condensate formation, but also a lack of condensate formation can cause human disease, underpinning the critical role of compartmentalization through phase separation for normal cell function and highlighting the emerging role of phase separation in the regulation of innate immune responses (Du and Chen, 2018; Hu et al., 2019; Zhou et al., 2021). Notably, AGS is primarily characterized by inflammation and neurodegeneration, raising the possibility that impaired formation of neuronal membraneless compartments might contribute to disease pathology and lending further support to the important role of inflammatory processes in neurodegeneration.

In summary, our findings define the regulation of cellular RNA concentrations by SAMHD1 as a key parameter for membraneless organelle assembly and delineate a novel mechanism by which compartmentalization of RNA in condensates buffers against erroneous immune recognition of self RNA.

## Acknowledgments

We thank Baek Kim, Martin Schlee, Georg Kochs, Kozo Kaibuchi, and Lucas Pelkmans for reagents, Vanessa Gilly and Kristin Eismann for technical assistance, and acknowledge the assistance of the TP Molecular Analysis / Mass Spectrometry, Center for Molecular and Cellular Bioengineering (CMCB), TU Dresden, and the imaging facilities of the CMBC, TU Dresden, and the Max Planck Institute of Molecular Cell Biology and Genetics (MPI-CBG), Dresden, the protein expression, purification and characterization facility, MPI-CBG, and the next generation sequencing platforms of the Max Delbrück Center for Molecular Medicine, Berlin, and the UMS2008 IBSLor, CNRS-Inserm-UL, Lorraine. This work was supported by grants from the Deutsche Forschungsgemeinschaft (KFO249 160548243 and CRC237 369799452/B21 to M.L.-K; CRC237 369799452/A01 and excellence cluster EXC2151 ImmunoSensation2 to G.H.). S.A. is supported by an European Research Council Consolidator Grant (725836). N.H is supported by an European Research Council Advanced Grant (AdG788970), the Leducq Fondation (16CVD03), the Chan Zuckerberg Initiative (2019-202666), and a BHF/DZHK grant (SP/19/1/34461). Y.M. was supported by FRCR project Epi-ARN from Grand Est Region, France. S.M. was supported by a fellowship of the Humboldt foundation (3.5-INI/1155756 STP). S.K. is recipient of a MeDDrive grant (60438) by the Medical Faculty, TU Dresden. N.C. is supported by SNSF_Early Postdoc Mobility-Stipendium (Nr. P2ZHP3_187653). The TP Molecular Analysis / Mass Spectrometry (CMBC) was supported by grants from the German Federal Ministry of Education and Research (BMBF) program ’Unternehmen Region’ (grants # 03Z2ES1, # 03Z22EB1), the German Research Foundation (INST 269/731-1 FUGG) and the European Regional Development Fund (ERDF/EFRE) (Contract # 100232736).

## Author contributions

S.K., S.M. and M. L.-K. conceived the project and designed experiments. S.A. contributed to the conceptual development. S.K. and S.M. performed experiments with help by S.H., N.C., N.L., and A.R.. X.Y., D.K. and A.H. performed *in vitro* experiments on condensate dissolution by ssRNA and dsRNA. S.T. and M.G. performed mass spectrometric analysis. T.Z., K.M. and G.H. generated genome-edited HeLa cells and nucleic acid ligands. V.M. and Y.M. performed RiboMethSeq analysis. H.M. and N.H. performed PAR-CLIP analysis. Y-T.C. contributed the F22 and E36 dyes. All authors contributed to data analysis and interpretation. The paper was written by S.M., S.K. and M. L.-K. with input from all authors.

## Declaration of interests

Simon Alberti and Anthony Hyman are advisors of Dewpoint Therapeutics. All other authors declare no competing interests.

## Methods

### Cell culture, transfection, cloning and drug treatment

Primary human fibroblasts, HeLa cells and HEK293 were grown in high glucose DMEM (ThermoFisher, 11995-065; Biochrom, F0435) containing 2 mM L-glutamine (ThermoFisher, 25030-024), 10% FCS (ThermoFisher, 26140-079; Biochrom, S0115) and 1X antibiotics/antimycotics (ThermoFisher, 10378-016). Fibroblasts from AGS patients, AGS1 (compound heterozygous for R290H and Q548X) and AGS2 (homozygous for H167Y) were derived from skin biopsies as previously described (Kretschmer et al., 2015). Plasmids or poly(I:C) (Invivogen, tlrl-pic) were transfected using either Lipofectamine 2000 (ThermoFisher, 11668-019) for HeLa cells or Viromer Red (Biontech, VR-01LB-01) for fibroblasts. For RNAi, HeLa cells were transfected using Oligofectamine (ThermoFisher, 12252-011) or Lipofectamine RNAiMAX (ThermoFisher, 13778-075) with siRNAs against SAMHD1 (sense RNA 5’-GCAGAUAAGUGAACGAGAUTT-3’, antisense RNA 5’-AUCUCGUUCACUUAUCUGCAG-3’) or the negative control #1 siRNA (Ambion). Three days after transfection with either siRNA or shRNA plasmids, cells were processed for further analysis. The SAMHD1-shRNA and the non-human shRNA-control plasmids were generated by cloning of oligonucleotides 5’-GCAGAUAAGUGAACGAGAUTT-3’ or 5’-GCAACAAGATGAAGAGCACCAA-3’, respectively, into pSicoR-mCherry vector (kind gift by Kozo Kaibuchi) (Funahashi et al., 2013). For shRNA-mediated knockdown of EXOSC10, RNASET2 or XRN1, HeLa cells were transfected using Lipofectamine2000 and 200 ng of MISSION shRNA plasmids (Sigma-Aldrich, TRCN0000006342, TRCN0000049850, and TRCN0000049677) or non-human control-shRNA. The next day, the medium was replaced with DMEM supplemented with puromycin (Invivogen, ant-pr-1) at a final concentration of 1 µg/ml. The medium was replaced every day to select for transfected clones. After 10-14 days, colonies were trypsinized and plated on coverslips in a 24-well plate for immunofluorescence staining or on 6-well plates for RT-PCR analysis.

Wild type and mutant human SAMHD1-cDNAs were cloned into pEGFP-C1 (GFP-SAMHD1) as previously described (Kretschmer et al., 2015). For induction of stress granules, cells were treated with 2 mM sodium arsenite (Sigma-Aldrich, 7784-46-5) for 1 h and for inhibition of stress granule formation cells were treated with 1.5 µg/ml of ISRIB (Sigma-Aldrich, SML0843). Stable HeLa cell lines constitutively expressing human marker proteins for membraneless compartments, SRRM1-GFP, G3BP1-mCherry, and nucleolin-GFP, were generated using BAC technology (Poser et al., 2008). HeLa BAC cell lines were kept under selection using geneticin (Gibco, 250 µg/ml).

Transient expression of GFP-DYRK3 was achieved by transfection of TH1668-(pOCC247)-pTT6-mGFP-DYRK3, created by cloning DYRK3 from pDONR223-DYRK3, which was a kind gift from William Hahn and David Root (Addgene, 23635) (Johannessen et al., 2010). For inducible GFP-DYRK3 expression, we used a HeLa-FlpIn-Trex cell line expressing GFP-tagged wild type DYRK3 (Rai et al., 2018), which was a kind gift from Lucas Pelkmans. Cells were maintained under selection with blasticidin (5 µg/ml) and hygromycin (200 µg/ml). DYRK3 expression was induced with 500 ng/ml of doxycycline for 1 h, after which cells were washed and fresh medium was added. Cells were further processed at the indicated times of incubation.

### Metabolic labeling of nascent RNA

Fibroblasts were grown on glass coverslips in DMEM containing non-essential amino acids, 2 mM L-glutamine, 10% FCS and 1X antibiotics/antimycotics. After 24 h, cells were pulse-labeled with the alkyne-modified nucleoside, 5-ethynyl-uridine (EU), at a final concentration of 1 mM for 30 min. Medium was then replaced with a fresh complete medium containing 2 µM actinomycin D (ThermoFisher, 11805017) to stop further transcription. At the indicated time points, cells were rinsed with PBS, fixed with 4% paraformaldehyde in PBS and processed for chemoselective ligation reaction between EU and azide using the Click-iT RNA Alexa Fluor 488 Imaging Kit (Molecular Probes, C10329). Finally, cells were mounted with VectaShield containing DAPI (Vector Laboratories, H-1200). Fluorescence images were captured using a LSM780 confocal microscope (Zeiss). For quantification of EU-AF488-labeled RNA, intensity after 30 min or *t* = 0 of EU labeling was normalized to 1.

### *In situ* RNA staining and quantification

HeLa cells or human fibroblasts were grown on glass coverslips in 24-well plates. After two quick washes with PBS, cells were fixed with 4% paraformaldehyde for 10 min and washed again with PBS. The cells were then stained for 10 minutes with 1 µM of the RNA dyes F22 or E36 (Li et al., 2006) dissolved in PBS and mounted with VectaShield containing DAPI (Vector Laboratory, H-1200-10) for quantification of total cellular RNA. Cells were imaged within 30 minutes of staining using a DeltaVision widefield microscope (GE) with a 60X objective.

dsRNA was immunostained using antibody J2 (SCICONS, 10010200) followed by Alexa Fluor 488-labeled secondary goat anti-Mouse IgG (Invitrogen, A-11029). Cells were imaged for dsRNA and DAPI at 60X magnification on a widefield microscope with an XY optical resolution of 0.1 µM X 0.1 µM and a Z step size of 0.35 µM or imaged with a 100X objective at the optical resolution 0.05 µM and a Z step size of 0.15 µM on a point scanning confocal microscope. Z-projected images of dsRNA were used to manually draw an outline of the cell boundary and generate a mask. This mask was then used on the best focused plane of the nucleus for automatic detection of the nuclear region using the DAPI channel. The same Z slice as the best focus plane of the nucleus was also used for quantification of the mean dsRNA intensity. Within the cytoplasm, a large fraction of focal dsRNA is localized in mitochondria (Dhir et al., 2018) as shown by co-staining with mitotracker red (Invitrogen, M7512). As dsRNA pattern recognition receptors are present in the cytoplasm, mitochondrial dsRNA foci were thresholded out by creating a mask for high intensity dsRNA foci. The remaining cytoplasmic region was then used for quantification of the mean dsRNA intensity.

For quantification of total RNA in fibroblasts, fractionation of nuclear and cytoplasmic fractions was carried out using the Ambion® PARIS™ kit (Invitrogen, Cat # AM1921). Total RNA was isolated with TRIzol™ (Invitrogen, Cat # 15596026) and quantified by spectrophotometric absorbance measurement at 260 nm.

### Immunofluorescence

Cells were fixed with 4% formaldehyde, permeabilized with 0.1% Triton X-100 in PBS for 10 min at room temperature and blocked with 2% BSA in PBS for 2 h at 4 °C before overnight incubation with the appropriate primary antibodies diluted in 2% BSA containing PBS at 4 °C. For nuclease treatment before antibody staining, fixed and permeabilized cells were incubated with PBS containing 10 µg/ml RNase A (ThermoFisher, EN0531) for 15 min in a humid chamber at 37 °C. After three washes with PBS, cells were incubated with appropriate Alexa Fluor-labeled secondary antibodies (Invitrogen, A-11029, A-32731, A-11035, A-11030) diluted in 2% BSA containing PBS for 2 h in the dark at room temperature. Finally, cells were washed for an additional three times with PBS before mounting with VectaShield containing DAPI. Fluorescence images were captured using a LSM780 confocal microscope (Zeiss) or DeltaVision widefield microscope (GE). The following antibodies were used: anti-DDX21 (GeneTex, GTX115199), anti-fibrillarin (Invitrogen, MA3-16771), anti-G3BP1 (ThermoFisher, PA5-29455), anti-dsRNA J2 (SCICONS, 10010200), anti-phosphorylated RNA pol II (Abcam, ab5131), anti-PML (Abcam, ab179466), anti-eIF3η (Santa Cruz, sc-137214) and anti-SC35 (Abcam, ab11826).

For quantification of SC35 nuclear speckles in fibroblasts, image stacks were recorded using an Operetta (Perkin Elmer) equipped with a 40X NA 0.95. For quantification (Perkin Elmer Harmony software) the following analysis pipeline was applied: Images were maximum projected and a flat field correction was applied. Nuclei were segmented: “Find Nuclei” method B (split factor 5, guide area 30 µm^2^, threshold 0.4, minimum contrast 0.1, split factor 5 and individual threshold 0.4). Nuclei were selected by the following parameters: Haralick contrast <0.8, 100 pixels < area 500 pixels, roundness >0.7 and intensity CD DAPI <30%. SC35 speckles were identified within nuclei using the “Find Spots” module A (Relative spot intensity >0.4, splitting coefficient 0.9) in the Alexa Fluor 488 channel.

For all other membraneless compartments, images were acquired at 60X magnification on a widefield microscope with an XY optical resolution of 0.1 µm X 0.1 µm and Z step size of 0.35 µM. Using a custom written Matlab routine, the cellular boundaries were marked by identification of the nuclear region using the DAPI staining and the relevant compartment was identified and quantified using the marker protein.

### RNA microinjection and RNA degradation assay

HeLa cells treated with control-shRNA or SAMHD1-shRNA for 3 days were plated on 35 mm coverslip bottomed plates (MatTek, P35G-1.5-14-C) one day before microinjection. The plates were mounted in a heating chamber on a microscope equipped with a micromanipulator and EMCCD camera to enable fast imaging. Cells were incubated with DMEM containing 10 mM HEPES for performing CO_2_-free microinjection and imaged at 37 °C. Microinjection was carried out using pre-pulled glass capillaries (Eppendorf Femtotips, 5242952.008) and the indicated RNA oligonucleotide diluted in nuclease-free water or just nuclease-free water using an Eppendorf GELoader (Eppendorf, 0030001222). The micromanipulator (Eppendorf InjectMan NI 2) was mounted on an Andor Spinning Disc Microscope and microinjection was performed using the microinjector FemtoJet with a 100X oil immersion objective to facilitate immediate visualization and image acquisition after microinjection. 50 hPa of compensation pressure and 120 hPA injection pressure (Pi) for 0.3 second injection time (Ti) was used for a single microinjection and a single image of the cells at the best focused plane was imaged before and after microinjection.

For microinjection experiments in HeLa cells stably expressing G3BP1-mCherry, SRRM1-GFP or nucleolin-GFP using ssRNA and dsRNA, the sequences of RNA oligonucleotides were as follows: PrD sense: 5’ AUU GAG GAG CAG CAG AGA AGU UGG AGU GAA GGC AGA GAG GGG UUA 3’ and PrD antisense: 5’ UAA CCC CUC UCU GCC UUC ACU CCA ACU UCU CUG CUG CUC CUC AAU 3’. dsRNA oligonucleotides were obtained by heating of complementary ssRNA oligonucleotides to 95 °C followed by slow cooling. Complete annealing of dsRNA oligonucleotides was verified by urea polyacrylamide gel electrophoresis. Single images at the best focused plane were imaged before and 10 min after microinjection. For quantification of condensate perturbation, the mean fluorescence intensity of the corresponding marker (G3BP1 for stress granules, SRRM1 for nuclear speckles and nucleolin for nucleoli) was quantified after microinjection.

To measure ssRNA degradation in cellulo, a 22 nt ssRNA probe labeled with 5’ FAM and 3’ TAMRA (5’-FAM-ACG CUG ACC CUG AAG UUC AUC -TAMRA-3’) was injected into the nucleus of HeLa cells treated with control-shRNA or SAMHD1-shRNA. The close proximity of TAMRA to FAM quenches the emission of FAM which can be excited at 488 nm wavelength excitation. Degradation of the probe releases the TAMRA quencher increasing FAM emission. Cells were imaged 10 min after microinjection and FAM fluorescence was recorded over the next 150 min during which fluorescence decreased due to photobleaching. The photobleaching rates varied due to different rates of ssRNA probe degradation which increases FAM fluorescence. To determine the rate of RNA probe degradation, the first image of FAM fluorescence intensity measured after 10 min of microinjection was normalized to 1 and the further drop in intensity was quantified. The RNA probe degradation index (D_RNA_) was calculated as follows:

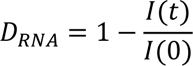

Where *I(t)* is the fluorescence intensity at time t and *I(0)* is the fluorescence intensity at time *t = 0*. The degradation curves were fitted using a single exponential function.

### Liquid chromatography-mass spectrometry for detecting nucleosides

Post-transcriptional modifications in RNA were determined by liquid chromatography/electrospray ionization mass spectrometry (LC-ESI-MS) of nucleoside digests. HeLa cells were grown in 6-well plates and transfected with SAMHD1-siRNA or control-siRNA. After 48 h, cells were washed twice with PBS, trypsinized and RNA was extracted using the RNeasy Mini Kit (Qiagen, 74104). Eluted RNA was incubated with 180 units of nuclease P1 (NEB, M0660) in nuclease P1 buffer at 37 °C for 90 min. The nuclease-digested RNA was then dephosphorylated by adding 10 units/µg of Quick CIP enzyme (NEB, M0525) and incubating at 37 °C for 90 min (Kellner et al., 2014). LC-MS/MS analysis was performed on a high-performance liquid chromatography system (Agilent 1200 HPLC) coupled to a Xevo G2-S QTof (Waters). For normal phase chromatography XBridge Amide 3.5 μm (2.1 x 100 mm) columns (Waters, 186004869) and for reverse phase chromatography CORTECS C18, 2.7 μm (2.1 x 100 mm) columns (Waters, 186007442) were used. The mobile phase was composed of 95% ACN, 0.1 mM NH_4_Ac and 0.01% NH_4_OH (eluent A) and 40% ACN, 0.1 mM NH_4_Ac and 0.01% NH_4_OH (eluent B). The following gradient program was used: Eluent B from 0% to 100% within 18 min; 100% from 18 min to 21min; 0% from 21 min to 26 min. The flow rate was set at 0.3 ml/min. The spray voltage was set at 3.0 kV and the source temperature was set at 120 °C. Nitrogen was used as both cone gas (50 l/h) and desolvation gas (800 l/h), and argon as the collision gas. MS^E^ mode was used with XBridge Amide 3.5 μm column in ESI negative ionization polarity for the detection of the nucleosides and positive ionization polarity with CORTECS C18, 2.7 μm column for the detection of the derivatized pseudouridine. Mass chromatograms and mass spectral data were acquired and processed by MassLynx software (Waters).

To identify pseudouridine in a mixture of nucleosides, derivatization with dansyl chloride was used. The dansyl chloride adducts attach to the N-1 position of the base in guanosine and guanosine-like residues and to the N-3 position in uridine and uridine-like residues. However, in the case of pseudouridine, the dansyl chloride adduct is substituted at both N-1 and N-3 positions of the base. To the volume of 100 µl of nucleoside mixture solution in H_2_O, 100 μl of 0.1 M HCl were added and incubated for 10 min at room temperature. Next, 100 μl of saturated Na_2_CO_3_ were added and incubated at room temperature for 1 h. Finally, 200 μl of 10 mM dansyl chloride solution in acetone were added and incubated at 4 °C overnight. To precipitate the extra dansyl chloride, 13 μl of 7% ammonia was added to the reaction tube and incubated at room temperature for 30 min. Then, the solution was centrifuged at 10,000 rpm for 5 min and 10 μl of the supernatant was directly injected into LC.

### Purification of recombinant proteins

Human SAMHD1 cloned into pGEX-6P-1 with an N-terminal GST tag (kind gift by Baek Kim) was purified as described (Amie et al., 2013). Briefly, transformed BL21 (DE3) competent cells (Novagen, 71397-4) were grown in 2X YT medium (Roth, X966.2) supplemented with 100 µg/ml ampicillin and 30 µg/ml chloramphenicol at 37 °C to an OD600 of 0.6, kept on ice for 2 h and induced with 0.25 mM IPTG (ThermoFisher, R1171) overnight at 25 °C. Cells were lysed in lysis buffer (50 mM Tris-HCl, pH 7.5, 500 mM NaCl, 2 mM EDTA, 1 mM DTT, 1 mg/ml lysozyme, one tablet of complete protease inhibitor cocktail (Roche, 04693124001) and one tablet of phosSTOP (Roche, 04906837001) for 4 h on ice. Cell debris was removed by centrifugation at 20,000 rpm for 20 min, and the resultant lysate was incubated overnight at 4 °C with 1.5 ml of glutathione-Sepharose 4B bead slurry (GE Healthcare, 17-0756-01). Beads were washed three times in wash buffer (50 mM Tris-HCl, pH 7.5, 500 mM NaCl, 1 mM DTT and 0.5% Triton X-100), equilibrated in buffer containing 50 mM Tris-HCl, pH 7.5, 150 mM NaCl, 10% glycerol and 0.5% Triton X-100, and packed into a polypropylene column (Qiagen, 34924). After two washes with 10 ml of equilibration buffer, GST-SAMHD1 was eluted with elution buffer (50 mM Tris-HCl, pH 8, 300 mM NaCl, 1 mM EDTA, 10% glycerol, 300 mM reduced glutathione). Eluted fractions were concentrated using Amicon Ultra-0.5 30K centrifugal filter devices (Merck Millipore, UFC5030BK) and desalted using protein desalting spin columns (ThermoFisher, 89849). Concentrated protein samples were separated on precast 12% polyacrylamide Bis-Tris gels (Invitrogen, NP0341BOX) and stained with Coomassie Blue to determine protein concentration by comparing them with a bovine serum albumin standard curve.

For SNAP-G3BP1 protein purification, baculoviruses were produced from a pOCC171-His_6_-MBP-3C-SNAP-GSAGSAAGSG-G3BP1 construct using an in-house-adapted SF9 insect cell expression system. Briefly, 500 ml SF9 cells (∼10^6^ cells/ml; Expression Systems, 94-001F) were infected with high-titer baculoviral supernatant, amplified in a second passage, and subsequently grown for 72 h at 27 °C and 85 rpm in ESF 921 culture medium (Expression Systems, 96-001-01). Cell pellets were obtained by centrifugation at 1000 rcf for 5 min and routinely flash frozen in liquid nitrogen (−196 °C) and then stored at -80 °C. For cell lysis, SF9 pellets were thawed and resuspended in 50 ml cold lysis buffer (50 mM Tris, 1 M NaCl, 20 mM imidazole, 5% (v/v) glycerol, 4 mM MgCl_2_, 1 mM DTT, 1x complete protease inhibitor, 1 U/ml DNase I, pH 7.4) and sonicated at 4 °C for overall 15 min, at an amplitude of 40% and throughout 30% of each cycle with an overall cycle length of 10 sec (Digital Sonifier S-450D, Branson). Following ultracentrifugation (17,000 rpm, 10 °C, 30 min), the obtained supernatant was passed through a 0.45 µm filter and subjected to a three-step FPLC purification on an ÄKTA pure 25 M chromatography system (Cytiva) at room temperature. Filtered cell lysates were first passed, at 5 ml/min, through the Ni^2+^ NTA resin of a preequilibrated 5 ml HisTrap HP column (Cytiva). This was followed by a washing step with 40 ml imidazole wash buffer (50 mM Tris, 100 mM NaCl, 20 mM imidazole, 5% (v/v) glycerol, pH 8), and an imidazole gradient elution over 50 ml, reaching 300 mM imidazole, under otherwise identical buffer conditions. Fractions, of 1.5 ml each, were collected throughout and those with high absorbance intensities at 280 nm were analyzed by SDS polyacrylamide electrophoresis, pooled, and passed through the quaternary ammonium cation-conjugated resin of a preequilibrated 5 ml HiTrap Q HP column (Cytiva). This was followed by a washing step with 40 ml Q wash buffer (50 mM Tris, 50 mM NaCl, 5% (v/v) glycerol, pH 8) and a high salt gradient elution over 50 ml, ultimately reaching 1 M NaCl and pH 7.4, under otherwise similar buffer conditions. Fractions, corresponding to the main eluting peak at 280 nm, were again analyzed and subsequently pooled. Following the concentration of pooled fractions, supplemented with additional 400 mM NaCl and 500 mM Arg-HCl (pH 7.4) to prevent aggregation, by repeated centrifugation (4,000 rcf, 25 °C, 3 min) in a 30 kDa cut-off filter column to ∼5 ml, purified proteins were digested with 500 µg 3C protease in presence of 0.5 mM DTT over 1 h at 25 °C. The resultant reaction volume was then further concentrated by centrifugation (4,000 rcf, 25 °C, 3 min) in a 30 kDa cut-off filter column to 1 ml. The 0.2 µm filtered concentrate was then resolved by size-exclusion chromatography (SEC) at a flow rate of 0.5 ml/min on a preequilibrated Superdex 200 Increase 10/300 GL column (∼24 ml column volume; Cytiva), in SEC buffer (50 mM Na_x_H_x_PO_4_, 300 mM NaCl, 5% (v/v) glycerol, 1 mM DTT, pH 7.4), which also constituted the final G3BP1 storage buffer. Following SEC, analysed and pooled fractions were again concentrated, as described above, in this case to 500 µl. Employing a ND-1000 spectrophotometer (ThermoFisher Scientific) the protein concentration was determined at 280 nm (with a molar extinction coefficient ε ∼39,880 M^-1^ cm^-1^ and a molecular weight of ∼72 kDa for SNAP-G3BP1 proteins) and the extent of nucleic acid contamination evaluated based on the 260 to 280 nm absorption ratio (obtaining a value of ∼0.62; a ratio of ≥0.7 indicating nucleic acid contamination). Subsequently, 8 µl aliquots were prepared, flash frozen in liquid nitrogen (−196 °C) and stored at -80 °C.

To exam the effect of dGTP on oligomerisation, purified GST-SAMHD1 (8 µg) was incubated with or without 200 µM dGTP in a total volume of 50 µl for 5 min prior to size exclusion chromatography (SEC) using an Ettan™ LC System (GE Healthcare). Samples were injected into an analytical Superdex 200PC3.2 column (GE Healthcare) with an approximate bed volume of 2.4 ml equilibrated with SEC buffer (20 mM Tris, 150 mM NaCl, 5 mM MgCl_2_, 0.5 mM DTT, pH 8). The absorbance was recorded at a wavelength of 280 nm. Following protein standards were used to estimate molecular weight: Thyroglobulin (660 kDa), Ferritin (440 kDa), Catalase (240 kDa) and Ovalbumin (43 kDa).

Phosphorylation of recombinant SAMHD1 proteins (15 µg) was carried out in a buffer containing 2 mM MOPS, 1 mM Glycerol-2-phosphat, 2 mM MgCl_2_, 400 µM EGTA, 400 µM EDTA, 20 µM DTT and phosSTOP in the presence of recombinant cyclin A2/cyclin-dependent kinase (CDK)1 (Sigma-Aldrich) and 10 mM ATP incubated for 2 h at 30 °C. Kinase reactions were subjected to immunoblotting with antibodies against phospho-SAMHD1 (T592) and SAMHD1 to validate phosphorylation.

### Mass spectrometry of recombinant SAMHD1 proteins

For in-solution digestion with trypsin and rLys-C protein, samples were diluted to 1 pmol/µl in 20 mM NH_4_HCO_3_. An aliquot of 50 µl from each sample was digested subsequently with trypsin (200 ng / 24 h) and rLys-C (100 ng / 24 h). Finally, 5 µl of the digested solution were acidified using 3 µl of 30% formic acid in water, supplemented with 25 fmol/µl standard peptide mixture (Pierce™ Peptide Retention Time Calibration Mixture, ThermoFisher, 88321) and diluted to a final volume of 23 µl with water. Five microliter of this solution were injected for nano LC-MS/MS. Nano LC-MS/MS analysis was performed on a Q Exactive™ HF hybrid quadrupole-Orbitrap mass spectrometer (ThermoFisher) hyphenated to a nanoflow LC system (Thermo Scientific™ Dionex UltiMate 3000 Rapid Separation LC (RSLC) system). Peptides were separated in a linear gradient of 0.1% aqueous formic acid (eluent A) and 0.1% formic acid in 60% acetonitrile (eluent B) over 45min, and the mass spectrometer operated in data-dependent acquisition mode (DDA, TopN 10). Raw files were converted to peak list files (mgf) with MSConverter (Kessner et al., 2008). Peptide and protein identification was performed with Mascot V2.6 (Matrixscience). In addition, peptide hits were evaluated and aggregated in Scaffold 4.11 (ProteomeSoftware) (Searle, 2010).

### *In vitro* nuclease assay

Purified recombinant SAMHD1 protein (3 µg) was added to reaction buffer containing 30 mM NaCl, 10 mM Tris-HCl (pH 8) and 2 mM CaCl_2_ in a volume of 25 μl and incubated with either 1 µg total RNA, 500 ng of poly(A)-RNA, 5 µg of 70S ribosome or 200 ng of synthetic RNA oligonucleotides, for 1 h at 37 °C. Benzonase (15 U; Merck Millipore, 70664-3) and nuclease-free water served as positive and negative control, respectively. After addition of 25 µl of 2X TBE-Urea sample buffer (Invitrogen, LC6876), reactions were heated to 70 °C for 2 min and 15 μl loaded on 15% polyacrylamide gels containing 7 M urea (Invitrogen, EC8852BOX). Low range ssRNA Ladder (NEB, N0364S) and dsRNA Ladder (NEB, N0363S) were used as size markers. Following electrophoresis, gels were stained by incubation in 50 ml ddH_2_0 containing 5 μl SYBR® Gold Nucleic Acid Gel Stain (Invitrogen, S11494) on a rocking platform for 30 min. Gels were imaged under UV light. Total RNA was extracted from HeLa cells using TRIzol (Ambion, 15596026) and stored at -80 °C. Mouse brain polyA+ RNA (ZYAGEN, MR-201_MR) and human brain microRNA (ZYAGEN, HR-201-SR) were purchased from Amsbio. The 3’-monophosphate oligonucleotides were purchased from BioSpring (3’p-ssRNA: 5’-UUAGGG-UUAGGGUUAGGGUUAGGGUUAGGGUUAG-p-3’). siRNA was purchased from ThermoFisher (5’p-CCGATGTGGAAGACCCCGAGGTGGAG-3’; dsRNA with 2 nt 3’ overhangs). 5’ppp dsRNA was purchased from Invivogen (tlrl-3prna). 5’-di- and triphosphate ssRNA oligonucleotides were synthesized as previously described (Goubau et al., 2014). Unmodified DNA oligonucleotides were synthesized by Eurofins MWG Operon and used for generation of RNA:DNA hybrids by annealing with the respective ssRNA oligonucleotides. The 5’-capped ssRNA oligonucleotide was kindly provided by Martin Schlee. All oligonucleotides were diluted in nuclease-free water to a concentration of 100 µM and stored at -80 °C. The *E. coli* 70S ribosome was purchased from NEB (P0763S).

To determine the effects of divalent cations on the RNase activity of SAMHD1, nuclease assays were performed in the presence of either 2 mM MgCl_2_, 200 pM ZnCl_2_, 2 mM CaCl_2_ and 200 pM ZnCl_2_, 2 mM MgCl_2_ and 200 pM ZnCl_2_ or a modified buffer containing 3 mM NaCl, 20 mM KCl, 10 mM Tris-HCl (pH 7.5) with 2 mM MgCl_2_, 200 pM ZnCl_2_ and 200 µM CaCl_2_. To exam the effect of oligomerisation of SAMHD1 on RNase activity, nuclease assays were performed in the presence of 200 μM dGTP.

### *In vitro* droplet assay

Stress granule-like *in vitro* condensates were prepared with modifications as described previously (Freibaum et al., 2021). Briefly, HeLa Kyoto cells were grown at 37 °C and 5% CO_2_ in DMEM high glucose (4.5 g/l) medium, supplemented with 10% FBS, 100 U/ml penicillin and 100 µg/ml streptomycin, to 70-80% confluency in 10 cm cell culture dishes. Cell pellets, routinely stored at -80 °C, were obtained by trypsinization (1 ml 0.05% trypsin, 3 min at 37 °C) of PBS-washed cells, followed by centrifugation at 1,000 rpm for 3 min, additional resuspension in 1 ml PBS, and a final centrifugation step at 4,000 rpm for 30 sec. To generate HeLa lysates, cell pellets were then thawed at room temperature and lysed by resuspension in 250 µl lysis buffer (50 mM Tris-HCl, 0.5% v/v Tergitol type NP-40, 0.2% v/v protease inhibitor cocktail III (Merck), 2.5% v/v murine RNase inhibitor (NEB), pH 7.0), followed by further incubation at room temperature for 3 min. Crude lysates were clarified by centrifugation at 21,000 g for 5 min and resultant supernatants were kept at 4 °C for several hours, during the course of the experiment.

Condensate formation was induced by addition of 20 µM SNAP-G3BP1, 5 mol% labelled with SNAP-Surface Alexa Fluor 488 (NEB, S9128S), to 25% v/v HeLa lysate in a final volume of 20 µl (50 mM Tris-HCl, pH 7.0). Reactions were supplemented with the indicated amounts of 45 nt PrD sense ssRNA (obtained from Sigma-Aldrich, 10 mol% 5’-Cy5-modified), 45 bp PrD dsRNA (PrD sense ssRNA annealed to PrD antisense ssRNA), rhodamine-labelled low molecular weight poly(I:C) (Invivogen, tlrl-piwr) or high molecular poly(I:C) (Invivogen, tlrl-picr) and condensate formation monitored at room temperature in 384-well µClear medium binding plates at indicated time points using an Eclipse Ti inverted spinning disc microscope (Nikon) with an iXON 897 EMCCD camera (Andor). Imaging was performed through a 60X 1.2 Plan Apochromat VC water-immersion objective (Nikon), using 488 nm and 660 nm lasers. The images were randomly acquired by selecting an automatic grid using the imaging software to reduce any bias. Images were then analyzed using a custom written Matlab routine. First, a background image was subtracted from the G3BP1 image to correct for effects of non-uniform illumination. Condensates were then thresholded using the G3BP1 intensity images such that any foci below 20 pixels in size were discarded. For a given image, the integrated G3BP1 density was computed for the protein in the condensate phase (G3BP1_cond_) and in the inverted mask made for the solution phase of the protein (G3BP1_sol_). The fraction of G3BP1 in the condensate was then computed as follows:

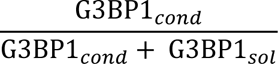

### Genome editing of HeLa cells

Hela cells were transiently electroporated with EF1alpha-Cas9-U6-sgRNA plasmid, using the Neon Transfection System (100 µg/ml plasmid, 1005 V, 35 ms, 2 pulses; ThermoFisher) and single-cell clones were obtained by limited dilution. CRISPR target sequences were designed with the Zhang lab CRISPR tool (crispr.mit.edu) or taken from the whole genome target list by Mali et al. (Mali et al., 2013). Gene-editing was confirmed by Sanger sequencing (Seqlab- Microsynth) and functional testing.

### Nucleic acid sensor mapping

For RNAi in primary fibroblasts, cells were transfected using Oligofectamine (ThermoFisher, 12252-011) with siRNAs against cGAS, MDA5, RIG-I, or the negative control #1 siRNA (all from Ambion). Three days after transfection, cells were processed for further analysis. For RNAi in HeLa cells, knockout cells (MDA5 ^-/-^; RIG-I ^-/-^, and MDA5^-/-^/RIG-I^-/-^ double knockouts) or wild type cells were transfected with Silencer Select small interfering RNAs (Ambion) targeting SAMHD1 (sense RNA 5’-GCAGAUAAGUGAACGAGAUTT-3’, antisense RNA 5’-AUCUCGUUCACUUAUCUGCAG-3’) or the negative control #1 siRNA using RNAiMAX (ThermoFisher, 13778-075). 24 h after transfection, cells were trypsinized and seeded into a 96-well plate at a density of 1 x 10^5^ cells per well. After additional 24 h of incubation, cells were stimulated with either 20 ng RIG-I ligand or 50 ng EMCV RNA combined with 200 U IFN-β (PBL, 11415-1) using 0.25 µl RNAiMAX per well. RIG-I agonist (5’ppp dsRNA) was generated by *in vitro* transcription with T7 RNA polymerase from a dsDNA template as previously described (Schlee et al., 2009). The reaction was then purified with TRIzol LS (ThermoFisher, 10296010) and Sephadex G-25 size-exclusion chromatography (GE, 17-0034-02). To generate MDA5-agonist (encephalomyocarditis virus [EMCV]-RNA), a confluent monolayer of Vero cells was infected with EMCV (Gitlin et al., 2006) at MOI of 1. After 12 h, total RNA was extracted with TRIzol. Activity and specificity were confirmed by transfection of wild type and MDA5-deficient murine bone marrow-derived dendritic cells.

Secretion of CXCL10 (IP-10) into the medium was quantified 24 h after stimulation by ELISA (Human IP-10; BD OptEIA, 550926). Briefly, ELISA plates were coated with 50 µl capture antibody at 4 °C overnight, washed three times with wash buffer (PBS, 0.05% Tween-20), blocked with assay buffer (10% FCS in PBS) for 1 h at room temperature, washed three times with wash buffer and incubated with either 50 µl standard or samples (diluted 1:2 in assay buffer) per well for 2 h at room temperature. Plates were then washed five times, incubated with 50 µl working detection mix (detection antibody and streptavidin-HRP) per well for 1 h and the reaction stopped by adding 25 μl of 1 M sulfuric acid. Plates were washed seven times before HRP activity was detected with TMB substrate solution in a microplate reader (Tecan, Infinite M200) at an absorbance of 450 nm.

### Quantitative RT-PCR

RNA was extracted with the ReliaPrep RNA Cell Miniprep System (Promega, Z6012) followed by DNase I digestion. RNA was reverse transcribed using the GoScript Reverse Transcription System (Promega, A5001). Target gene expression was determined by quantitative RT-PCR using GoTaq qPCR Master mix (Promega, A6002) on a QuantStudio 5 Real-Time PCR System (Applied Biosystems) and normalized to GAPDH. The following primers were used: CXCL10-fwd: 5’-GTGGCATTCAAGGAGTACCTC-3’ and CXCL10-rev: 5’-TGATGGCCTTCGATTCTGGATT-3’; MX1-fwd: 5’-GTTTCCGAAGTGGACATCGCA-3’ and MX1-rev: 5’-CTGCACAGGTTGTTCTCAGC-3’; GAPDH-fwd: 5’-GAAGGTGAAGGT-CGGAGTC-3’ and GAPDH-rev: 5’-GAAGATGGTGATGGGATTTC-3’. For determination of shRNA-mediated knockdown efficiency in HeLa cells, the following qPCR primers (Origene) were used: EXOSC10-fwd: 5’-CTC TTT GGA CCT CAC GAC TGC T-3’; EXOSC10-rev: 5’-AAG AGG CTC GCC TGC TTC TGA A-3’; XRN1-fwd: 5’-CCA GCA AAG CAG TCG TGG AGA A-3’; XRN1-rev: 5’-CCA CGA CTC TAG CTT CCT CAA G-3’; RNASET2-fwd: 5’-AAT CCA GTG CCT TCC ACC AAG C-3’, and RNASET2-rev: 5’-CCA TTT GCC AGC CAG ACT TCC T-3’.

### Western blot analysis

Cells were lysed in RIPA buffer (50 mM Tris-HCl, pH 7.4, 150 mM NaCl, 1 mM EDTA, 1% Triton X-100, 1 mM Na_3_VO_4_ and 20 mM NaF) supplemented with 2 U/ml DNase I (Invitrogen, AM2222), protease and phosphatase inhibitors. Proteins were separated by electrophoresis on 4-12% NuPAGE gels (Invitrogen, NP0321BOX, NP0322BOX) and analyzed by Western blotting using the following antibodies: anti-SAMHD1 (ProteinTech Group, 12586-1-AP), anti-phosphoSAMHD1 (T592) (ProSci, 8005), anti-cGAS (Cell Signaling, 15102S), anti-RIG-I (Cell Signaling, 3743S), anti-MDA5 (Cell Signaling, 5321S), anti-GFP (Roche, 11814460001), anti-MX1 (kindly provided by Georg Kochs), anti-pSTAT1 (Cell Signaling, 7649S), anti-STAT1 (Cell Signaling, 14994S) and anti-β-actin (Sigma Aldrich, AC-15). Immunoreactive signals were detected by chemiluminescence using Lumi-Light PLUS (Roche, 12015196001).

### RiboMethSeq analysis of rRNA 2’-O-methylation

Levels of 2’-O-methylations at 110 known sites in human rRNA were determined using the RiboMethSeq protocol (Marchand et al., 2016). In brief, total RNA was subjected to random alkaline hydrolysis followed by end-repair and ligation of 3’-end and 5’-end adapters to RNA fragments. The amplified library was sequenced on Illumina HiSeq 1000 in a single-read SR50 mode. All bioinformatic steps (trimming, alignment and counting) were performed essentially as described previously using +/-2 nt neighboring interval (Ayadi et al., 2018). Calibration of absolute 2’-O-Me level was performed using *in vitro* human rRNA transcripts covering all modified positions. Differential heatmap was constructed by hclust function and plotted by heatmap.2 of R package (v.3.5.3).

### Photoactivatable-ribonucleoside-enhanced crosslinking and immunoprecipitation

Photoactivatable-ribonucleoside-enhanced crosslinking and immunoprecipitation (PAR-CLIP) libraries were generated and analyzed as previously described (Hafner et al., 2010; Maatz et al., 2014). HEK293 cells stably expressing wild type (GFP-SAMHD1) or mutant SAMHD1 (GFP-SAMHD1_H167Y; GFP-SAMHD1_Q548X) were grown for 16 h in medium supplemented with 4-thiouridine (4SU, Sigma T4509) at a final concentration of 100 µM. After washing with PBS, cells were irradiated with 0.15 J/cm² of 365 nm UV light in a Stratalinker 2400 crosslinker (Stratagene) and harvested. Cell pellets were lysed in NP40 lysis buffer (50 mM HEPES, pH 7.5, 150 mM KCl, 2 mM EDTA, 1 mM NaF, 0.5% (v/v) NP40, 0.5 mM DTT, complete protease inhibitor cocktail) and the cleared cell lysates were treated with 1 U/µl RNase T1 (ThermoFisher, EN0541). GFP-tagged SAMHD1 proteins were immunoprecipitated with anti-SAMHD1 antibody (Sigma-Aldrich, SAB1400478) bound to Protein-A/G-Agarose (ThermoFisher, 20422) and subsequently treated with 100 U/µl RNase T1. Beads were washed and resuspended in NEBuffer 3 containing Calf Intestinal Alkaline Phosphatase at a concentration of 0.5 U/μl (NEB, M0290) to dephosphorylate the RNA. Beads were washed with phosphatase wash buffer (50 mM Tris-HCl, pH 7.5, 20 mM EGTA, 0.5% (v/v) NP40) followed by polynucleotide kinase (PNK) Buffer (50 mM Tris-HCl, pH 7.5, 50 mM NaCl, 10 mM MgCl_2_, 5 mM DTT) prior to incubating with 0.5 μCi/μl [γ^32^P]ATP and T4 PNK (NEB, M0201L) to radiolabel crosslinked RNA. The protein-RNA complexes were separated on 4-12% NuPAGE Midi gels (Invitrogen, WG1402BOX) and electroeluted. The electroeluate was digested with 2 mg/ml proteinase K for 1 h at 55 °C. The RNA was extracted using phenol/chloroform/isoamylalcohol (25:24:1 pH 7.4) followed by ethanol precipitation. The recovered RNA was converted into a PAR-CLIP cDNA library and sequenced as described previously (Hafner et al., 2010). The resulting SAMHD1 binding site datasets were mapped to the human transcriptome (Ensembl v.75) and the number of peaks overlapping annotated exons, 5’ UTRs, 3’ UTRs and introns quantified. UTR regions overlapping alternative CDS sequences and intronic regions overlapping alternative exonic sequences were excluded (Maatz et al., 2014).

### Statistical analysis

Comparison of means was carried out in GraphPad PRISM6 using Student’s t-test, Mann-Whitney-U test or ANOVA followed by post-hoc test as indicated. p*<*0.05 was considered significant. Data are presented as mean ± SD or mean ± SEM as indicated. In PRISM, D’Agostino-Pearson omnibus K2 test was used to determine whether the data met assumptions of the statistical approach.

## Supplemental Information

**Figure S1.**
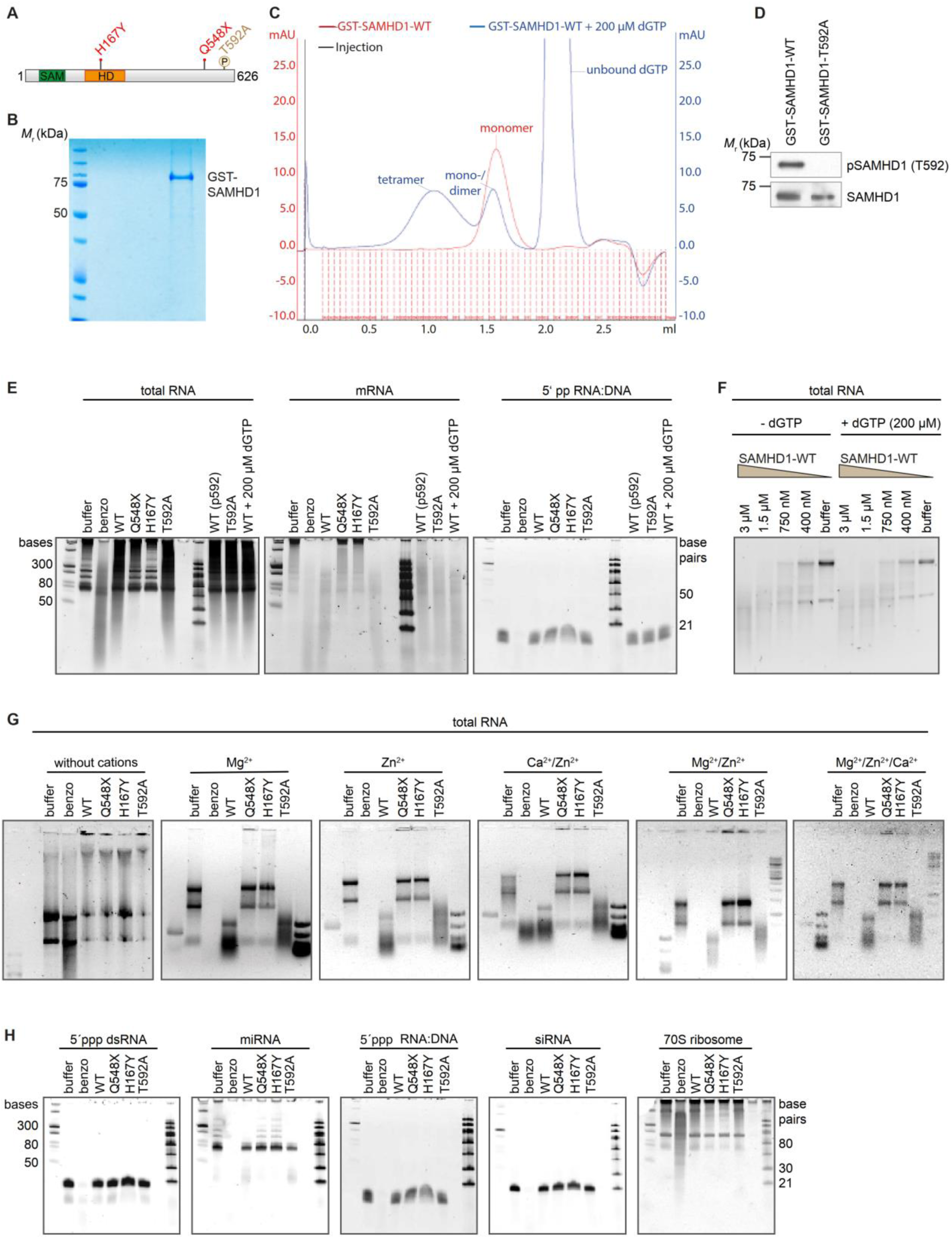
Characterization of RNase activity of SAMHD1 *in vitro* and *in cellulo*. (**A**) Schematic illustration of the human SAMHD1 protein and its functional domains (SAM domain, aa45-110 and catalytic HD domain, aa164-316). Investigated mutations (H167Y, Q548X, and T592A) are highlighted. (**B**) Coomassie Blue-stained SDS-PAGE of purified GST-SAMHD1. (**C**) Size exclusion chromatography profiles of GST-SAMHD1 alone (red) or in the presence of 200 µM dGTP (blue). (**D**) Western blot of wild type (GST-SAMHD1-WT) and phosphorylation-deficient SAMHD1 (GST-SAMHD1-T592A) after treatment with recombinant cyclin A2/cyclin-dependent kinase (CDK) 1, immunostained with antibody against SAMHD1 or phospho-SAMHD1 (T592). (**E**) Last three lanes of each panel: RNase activity of phosphorylated wild type SAMHD1 (WT p592) and phosphorylation-deficient SAMHD1 (T592A) after treatment with recombinant cyclin A2/CDK1, and wild type SAMHD1 in the presence of 200 µM dGTP (WT + 200 µM dGTP) using the following substrates: total RNA, mRNA and RNA:DNA hybrid with 5’ diphosphate on RNA (5’ pp RNA:DNA). Buffer, negative control. Benzo, benzonase, positive control. First lane, ssRNA ladder; last lane, dsRNA ladder. (**F**) RNase activity of wild type SAMHD1 without or in the presence of 200 µM dGTP using total RNA as substrate. (**G**) RNase activity of wild type SAMHD1 without or in the presence of the indicated divalent cations (either 2 mM Mg^2+^, 200 pM Zn^2+^, 2 mM Ca^2+^ and 200 pM Zn^2+^, 2 mM Mg^2+^ and 200 pM Zn^2+^ or a modified buffer containing 3 mM NaCl, 20 mM KCl, 10 mM Tris-HCl (pH 7.5) and 2 mM Mg^2+^, 200 pM Zn^2+^ and 200 µM Ca^2+^) using total RNA as substrate. (**H**) RNase activity of wild type (WT) and mutant (Q548X, H167Y, T592A) SAMHD1 using the following substrates: dsRNA carrying 5’ triphosphate (5’ppp dsRNA), primary micro-RNA (miRNA), RNA:DNA hybrid with 5’ triphosphate on RNA (5’ ppp RNA:DNA), small interfering RNA (siRNA) and yeast 70S ribosome. Buffer, negative control. Benzo, benzonase, positive control. First lane, ssRNA ladder; last lane, dsRNA ladder.

**Figure S2.**
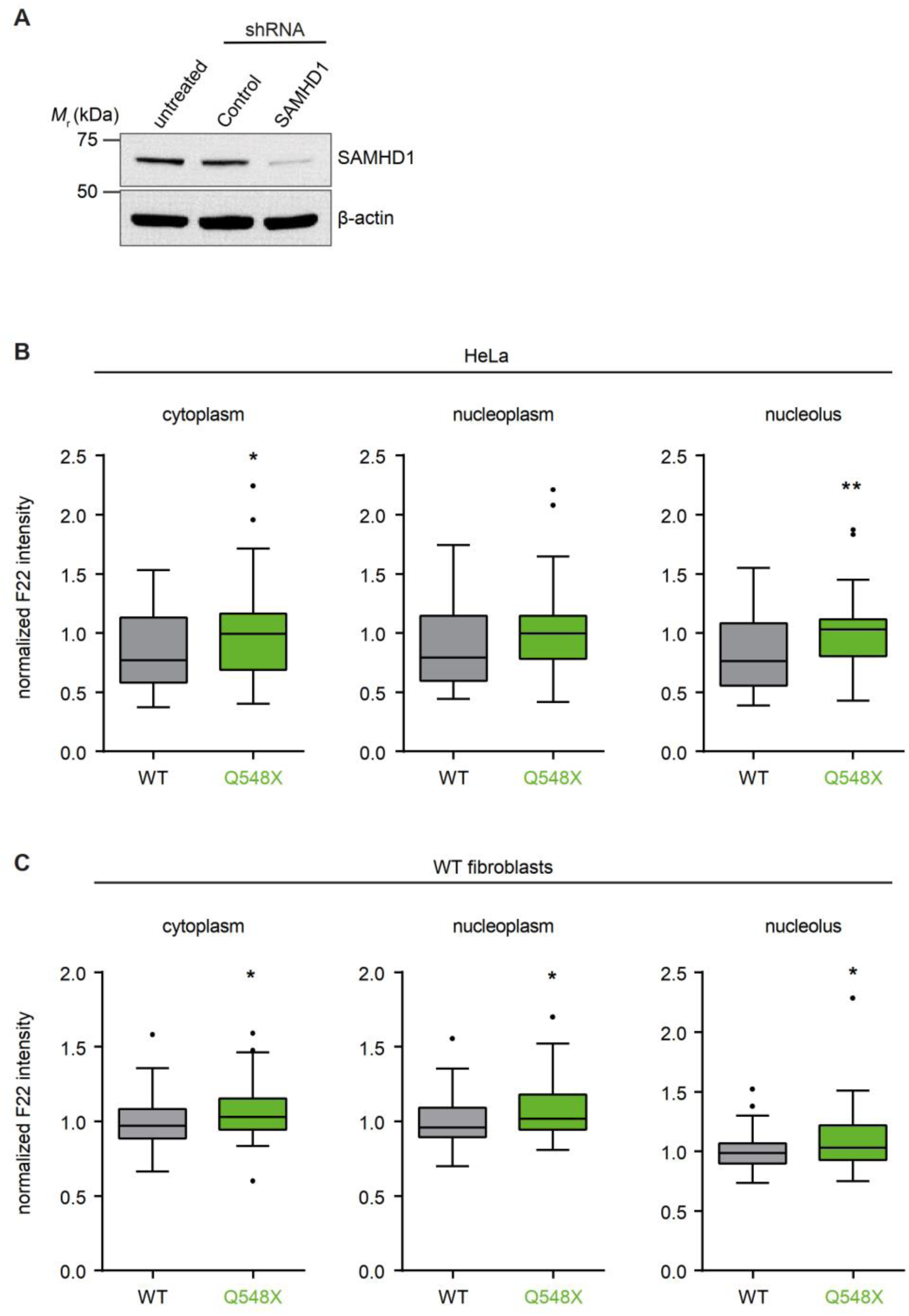
SAMHD1 depletion causes cellular accumulation of RNA. (**A**) Immunoblot of HeLa cells treated with control-shRNA or SAMHD1-shRNA using indicated antibodies. (**B**, **C**) Quantification of RNA stained with F22 dye in HeLa cells (n = 52-60, **B**) and wild type fibroblasts (n = 40-61, **C**) overexpressing wild type or mutant (Q548X) SAMHD1. Data are representative of at least three independent experiments. Box plots: center line, median; box, interquartile range; whiskers, 1.5x interquartile range; dots, outliers. Mann-Whitney U test. *, p<0.05; **, p<0.01.

**Figure S3.**
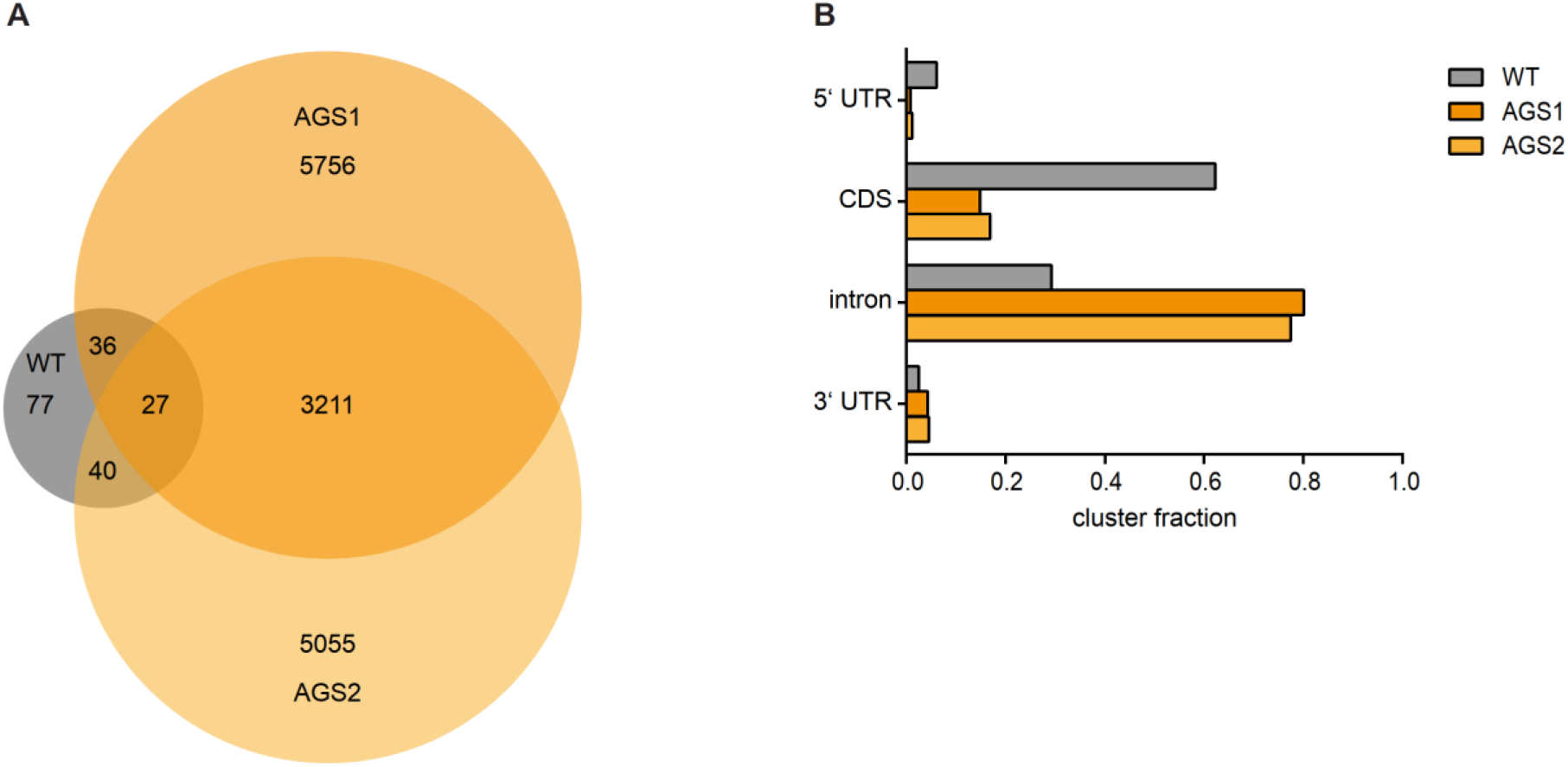
PAR-CLIP analysis for quantification of RNA bound to SAMHD1. (**A**) Venn diagram showing number of genes targeted by wild type (WT) and mutant SAMHD1 (AGS1, Q548X; AGS2, H167Y) in stably transfected HEK293 cells. (**B**) Distribution of sequence clusters across intronic and exonic regions of RefSeq mRNAs captured by PAR-CLIP.

**Figure S4.**
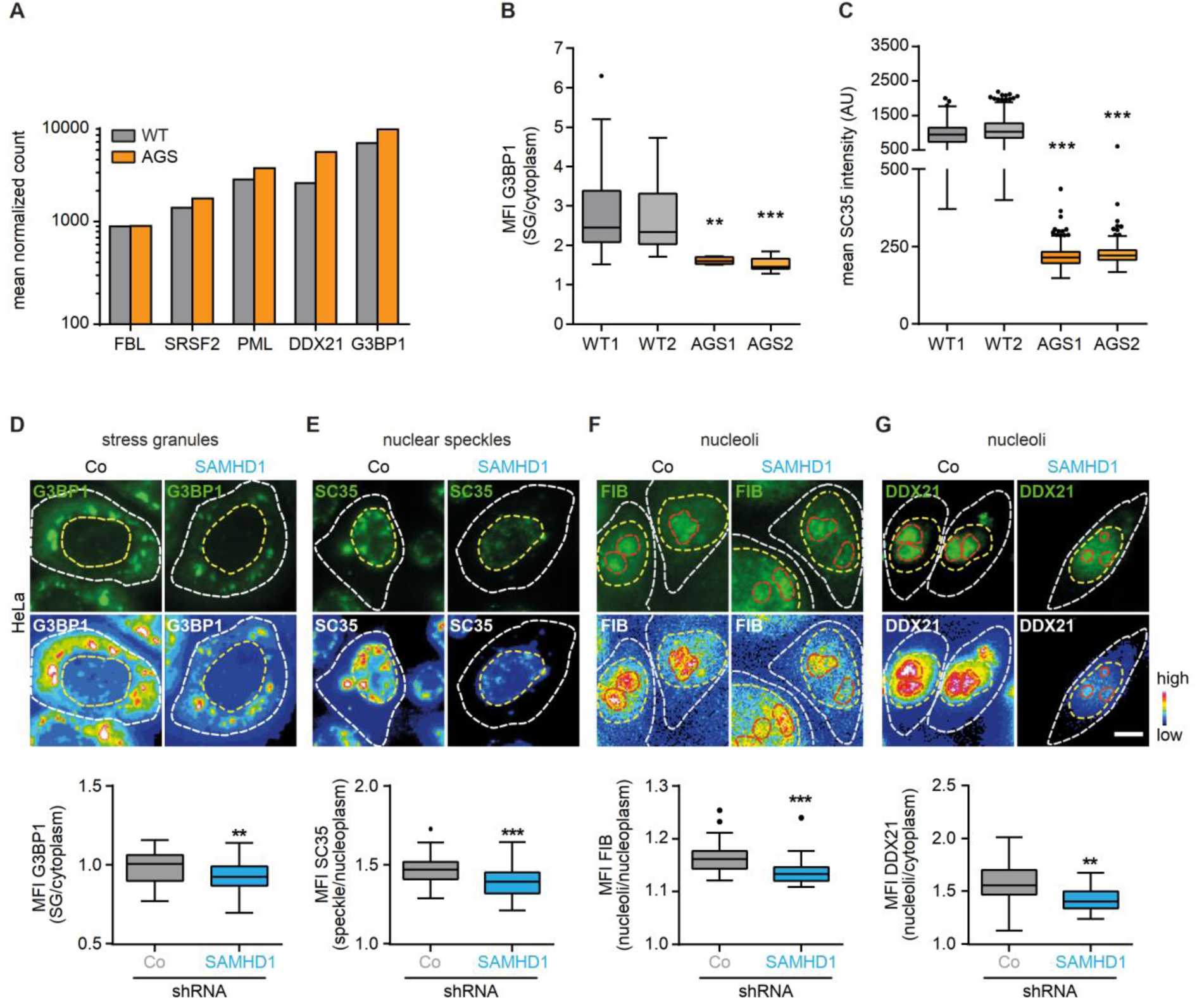
Structural perturbation of membraneless compartments in SAMHD1-deficient cells. (**A**) mRNA expression of condensate-specific constituent proteins in AGS patient cells and wild type controls. (**B**, **C**) Mean fluorescence intensity (MFI) of G3BP1 in stress granules relative to cytoplasm (n = 94-135 cells, **B**) and of SC35 in nuclear speckles (n = 222-559 cells, **C**). Box plots: center line, median; box, interquartile range; whiskers, 1.5x interquartile range; dots, outliers. Kruskal-Wallis test with Dunn’s multiple comparisons test. **, p<0.01; ***, p<0.001. Data are representative of three independent experiments. (**D**-**G**) Images and quantification of membraneless compartments in HeLa cells transfected with control-shRNA (Co) or SAMHD1-shRNA (SAMHD1). White dotted lines mark the cellular boundary, and yellow dotted lines denote the nuclear outline. Scale bars, 5 μm. G3BP1, n = 41-89 cells; SC35, n = 61 cells; FIB, n = 56-60 cells; DDX21, n = 41-89 cells. Box plots: center line, median; box, interquartile range; whiskers, 1.5x interquartile range; dots, outliers. t-test. **, p<0.01; ***, p<0.001. Data are representative of three independent experiments.

**Figure S5.**
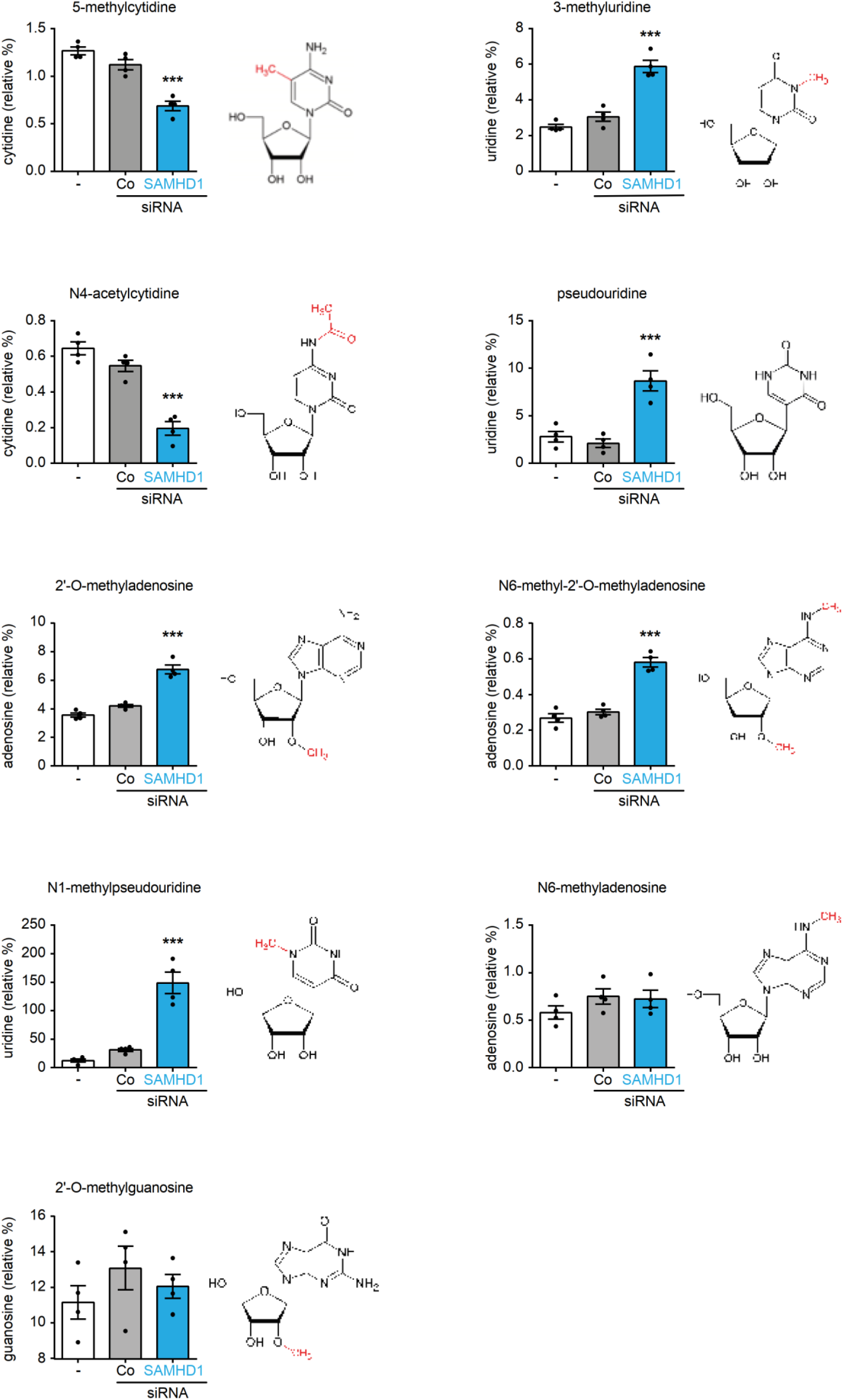
Functional perturbation of membraneless compartments in SAMHD1-deficient cells. Quantification of ribonucleoside modifications in untreated HeLa cells (-) and cells treated with control-siRNA (Co) or SAMHD1-siRNA (SAMHD1). One-way ANOVA, control-siRNA vs. SAMHD1-siRNA. ***, p<0.001. Mean ± SEM of triplicate technical replicates, representative of two independent experiments.

**Figure S6.**
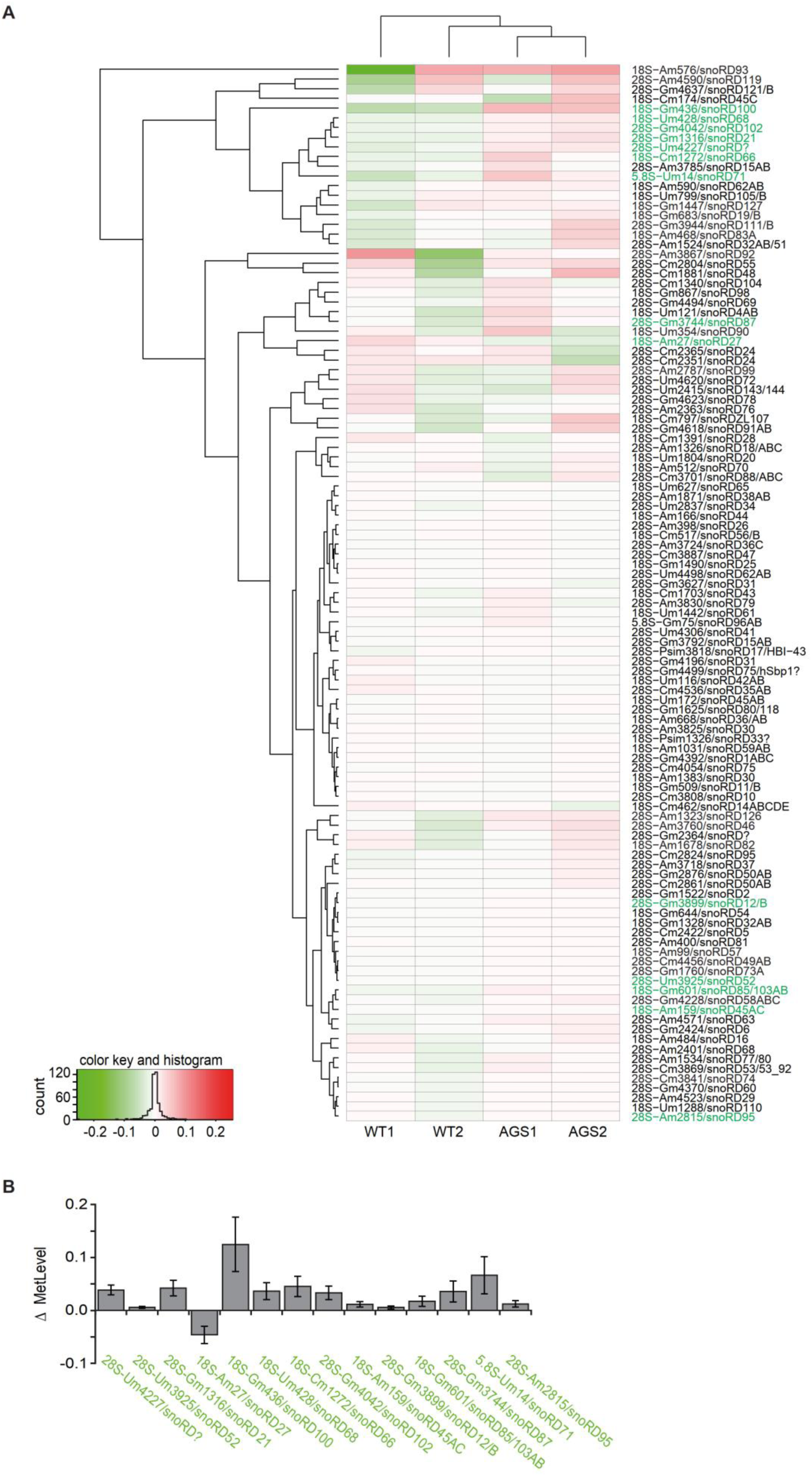
Altered 2’-O-methylation of rRNA in SAMHD1-deficient AGS patient fibroblasts. (**A**, **B**) Differential heatmap of rRNA methylation as constructed by site-by-site normalization of RNA methylation level (**A**). Red colors correspond to increased methylation compared to average, while green colors show undermethylated positions. Color key and histogram are shown on the bottom. Identity of rRNA positions includes the type of rRNA, modified nucleotide and its number, as well as the associated C/D-box snoRNA. Positions highlighted in green differ with p-value <0.05 and are also shown as differential plot between patients and controls (**B**). Y-axis represents the difference in methylation between patients and controls.

**Figure S7.**
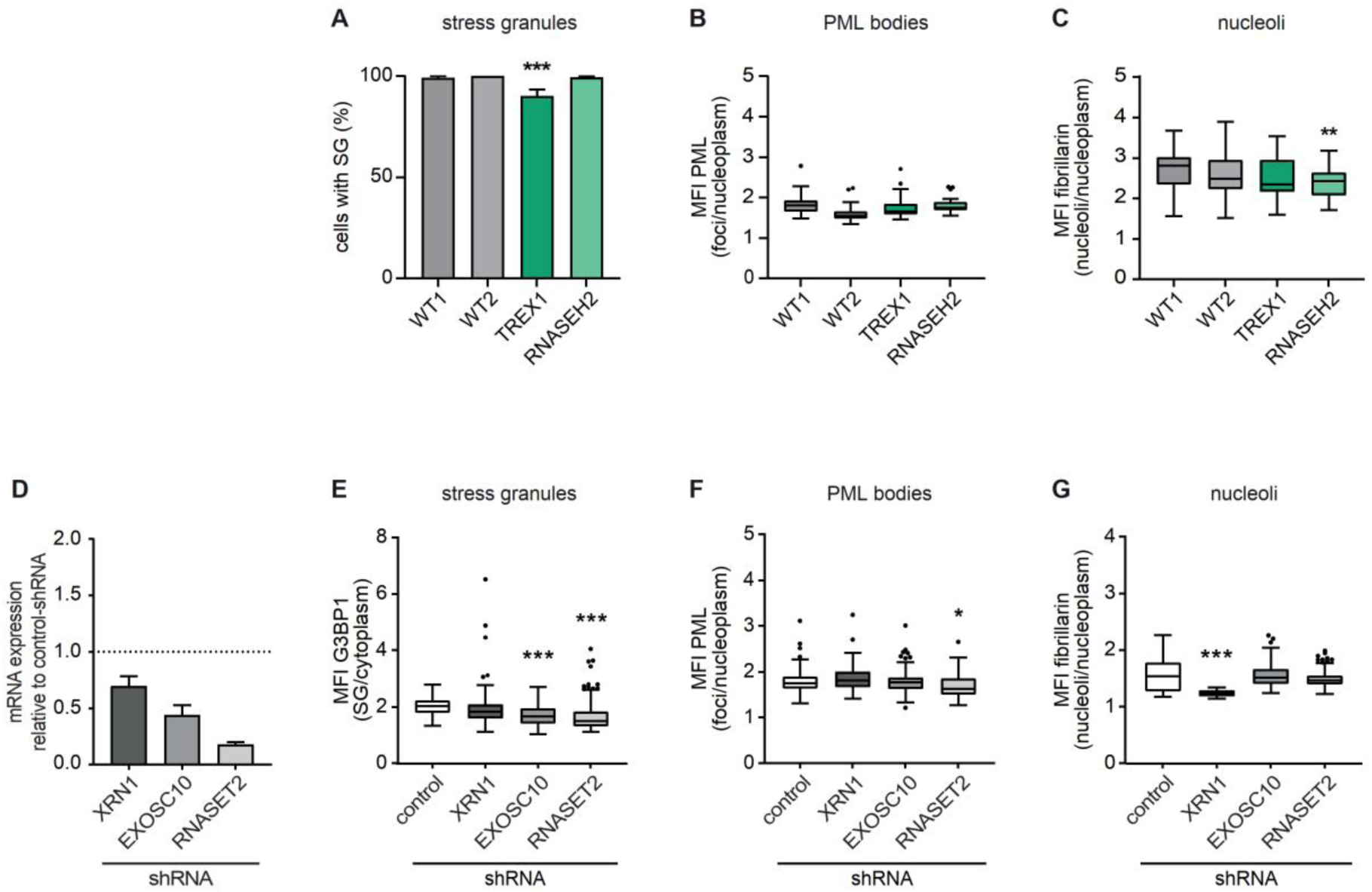
Lack of other cellular nucleases does not globally disturb condensate formation. (**A**-**C**) Quantification of stress granules, PML bodies, and nucleoli after staining of marker proteins, G3BP1, PML, and fibrillarin, respectively, in wild type (WT1, WT2) and AGS patient fibroblasts with mutations in TREX1 and RNASEH2. Stress granules are shown as percent of cells with stress granules, n = 20-26 cells. PML bodies, n = 25-41 cells. Nucleoli, n = 27-50 cells. Box plots: center line, median; box, interquartile range; whiskers, 1.5x interquartile range; dots, outliers. Kruskal-Wallis test with Dunn’s multiple comparisons test. ***, p<0.001. Data are representative of two independent experiments. (**D**) Validation of shRNA-mediated knockdown of XRN1, EXOSC10 and RNASET2 by quantitative RT-PCR. (**E-G**) Quantification of stress granules, PML bodies, and nucleoli in HeLa cells treated with shRNA against XRN1, EXOSC10, RNASET2, or non-human control after staining of marker proteins, G3BP1, PML, and fibrillarin, respectively. Stress granules are shown as enrichment of G3BP1 in stress granules, n = 57-185 cells. PML bodies, n = 44-157 cells. Nucleoli, n = 15-273 cells. Box plots: center line, median; box, interquartile range; whiskers, 1.5x interquartile range; dots, outliers. Kruskal-Wallis test with Dunn’s multiple comparisons test. ***, p<0.001. Data are representative of two independent experiments.

**Figure S8.**
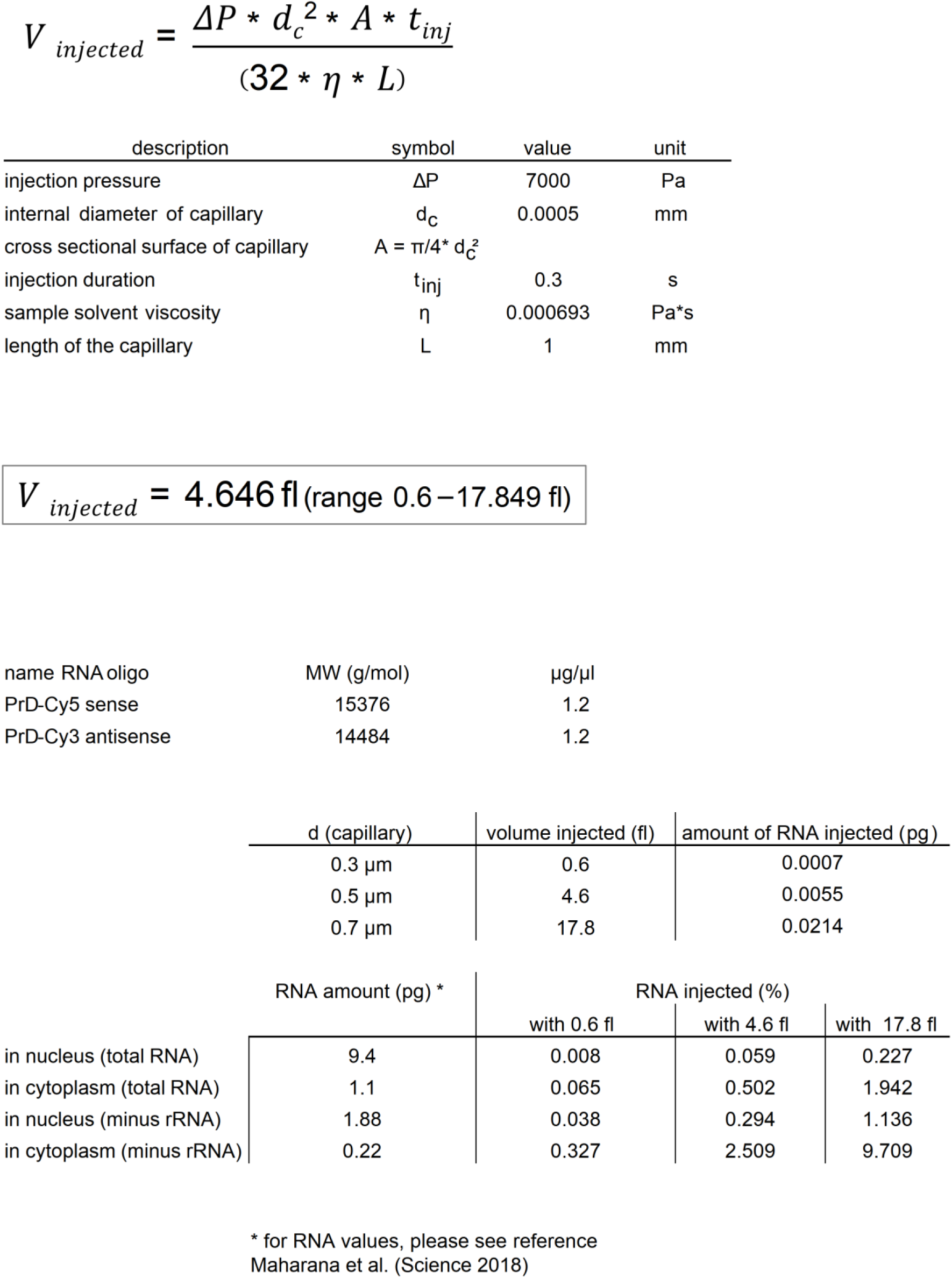
Estimation of RNA amounts injected into HeLa cells. The equation shows calculation of the volume microinjected into HeLa cells, where ΔP is the injection pressure, d_c_ is the inner diameter of the injection capillary (0.5 µm ± 0.2 µm), A is the cross sectional surface of the capillary, t_inj_ is the injection duration, η is the sample solvent viscosity and L is the length of the capillary. Because of the variable internal diameter of the capillaries, the calculated injected volume ranges between 0.6-17.8 fl. Given the concentration of the injected RNA of 1.2 µg/µl, the amount of injected RNA ranges between 0.0007-0.0214 pg. Taken into account that a HeLa cell contain on average 9.4 pg total RNA in the nucleus and 1.1 pg total RNA in the cytoplasm (Maharana et al., 2018), the injected amount of RNA corresponds to up to ∼0.2% of RNA in the nucleus and up to ∼2% in the cytoplasm. When subtracting the ribosomal RNA from the total RNA, the microinjected amount of RNA corresponds to up to ∼1.1% RNA in the nucleus and to up to ∼9.7% in the cytoplasm.

**Figure S9.**
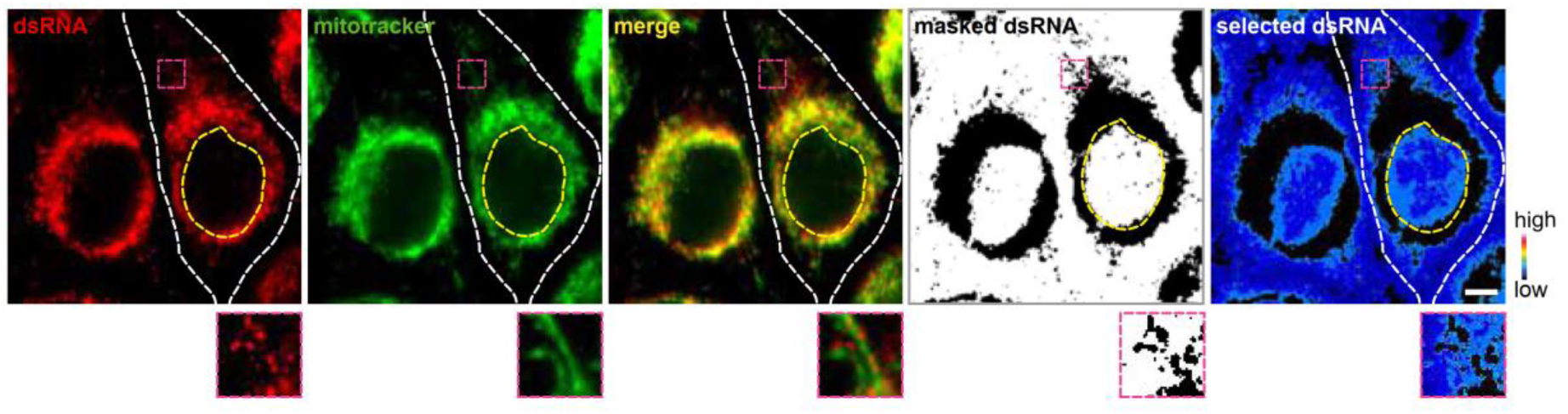
*In situ* immunostaining of dsRNA. J2 imaging mask. Representative images of HeLa cells stained with mitotracker (green) and immunostained for dsRNA (red). The black and white image shows the masked region of the dsRNA image after thresholding for dsRNA foci. Only the white region is selected for further quantification. The last image shows the color-coded intensity of dsRNA selected for analysis. Outlines of cells and nuclei are marked with white and yellow dashed lines, respectively. Scale bar, 5 μm.

**Table S1.** Mass spectrometry of purified recombinant SAMHD1. MS/MS samples were analyzed using Mascot (Matrix Science, London, UK; version 2.6.0). Mascot was set up to search the SwissProt_2020_05.fasta; Enzymes_MSG_20180829.fasta; STDs_TAGs_20200212.fasta; Contaminants_MSG_20170718 database (563701 entries) assuming the digestion enzyme trypsin. Scaffold (version Scaffold_4.8.4, Proteome Software Inc., Portland, OR) was used to validate MS/MS based peptide and protein identifications.

| Identified Proteins | Accession Number | Molecular Weight | Total Spectrum Count |
| --- | --- | --- | --- |
| Deoxynucleoside triphosphate triphosphohydrolase SAMHD1, Homo sapiens | Q9Y3Z3 | 72 kDa | 398 |
| Deoxynucleoside triphosphate triphosphohydrolase SAMHD1, Mus musculus | Q60710 | 76 kDa | 45 |
| Deoxynucleoside triphosphate triphosphohydrolase SAMHD1, Xenopus laevis | Q6INN8 | 73 kDa | 32 |
| 60 kDa chaperonin, Escherichia coli | A1AJ51 | 57 kDa | 88 |
| 60 kDa chaperonin OS=Citrobacter koseri | A8AMQ6 | 57 kDa | 71 |
| 60 kDa chaperonin OS=Haemophilus influenzae | A5U9V2 | 58 kDa | 42 |
| Glutathione S-transferase class-mu 26 kDa isozyme OS=Schistosoma japonicum | P08515 | 26 kDa | 65 |
| P00761 Trypsin OS=Sus scrofa | Z99999 | 24 kDa | 26 |
| Chaperone protein DnaK OS=Escherichia coli O1:K1 | A1A766 | 69 kDa | 18 |
| Protein RecA OS=Escherichia coli O139:H28 | A7ZQC8 | 38 kDa | 10 |
| ATP-dependent protease ATPase subunit HslU OS=Escherichia coli O1:K1 | A1AIA6 | 50 kDa | 6 |
| ATP-dependent protease ATPase subunit HslU OS=Buchnera aphidicola | Q89A31 | 51 kDa | 1 |
| Chaperone protein DnaJ OS=Escherichia coli O139:H28 | A7ZHA5 | 41 kDa | 6 |
| Albumin OS=Homo sapiens | P02768 | 69 kDa | 4 |
| Keratin, type II cytoskeletal 2 epidermal - Homo sapiens | K22E_HUMAN | 66 kDa | 3 |
| Elongation factor Tu 1 OS=Escherichia coli O1:K1 | A1AGM6 | 43 kDa | 3 |
| Keratin, type I cytoskeletal 10 - Homo sapiens | K1C10_HUMAN | 60 kDa | 2 |
| Keratin, type II cytoskeletal 1 OS=Pan troglodytes | A5A6M6 | 65 kDa | 2 |
| 30S ribosomal protein S5 OS=Escherichia coli O1:K1 | A1AGJ2 | 18 kDa | 2 |
| D-tagatose-1,6-bisphosphate aldolase subunit GatZ OS=Escherichia coli O139:H28 | A7ZNR4 | 47 kDa | 2 |
| ATP-dependent Clp protease ATP-binding subunit ClpX OS=Shewanella sp. | A0KYL8 | 47 kDa | 2 |

